# An orphan LuxR regulator controls plant developmental reprogramming during interaction with *Novosphingobium resinovorum* HR1a

**DOI:** 10.64898/2026.11.03.762470

**Authors:** Lázaro Molina, Irene Hurtado, Zulema Udaondo, Ana Segura

## Abstract

Rhizoremediation depends on productive plant/pollutant-degrading microorganism interactions, yet the bacterial regulatory mechanisms that shape host responses remain poorly understood. We investigated the role of the previously uncharacterized orphan LuxR874 from the polycyclic aromatic hydrocarbon (PAH)-degrading bacterium *Novosphingobium resinovorum* HR1a, in the modulation of plant physiology. Bacterial transcriptomics revealed that LuxR874 controls the expression of genes associated with aromatic and sugar metabolism, and acyl-CoA-related functions. Inactivation of *luxR874* altered bacterial responses without affecting root colonization, suggesting that this regulator contributes to the physiological state required to elicit host responses. In gnotobiotic systems, HR1a induced a root phenotype characterized by reduced primary root elongation, increased root thickness, and enhanced root hair formation. These alterations were attenuated in the *luxR874* mutant and restored by complementation despite similar colonization levels among strains. Plant RNA-seq analyses revealed extensive host transcriptional reprogramming, including activation of developmental regulators, hormone-associated pathways, cell-wall remodeling genes, and oxidative stress responses. In contrast, plants inoculated with the *luxR874* mutant displayed a weaker transcriptional response resembling the non-inoculated state. Functional analyses supported LuxR874-dependent activation of reactive oxygen species-and ethylene-related pathways. Our results reveal an unexpected role for an orphan LuxR regulator in coordinating multi-level responses during a rhizoremediation-relevant plant–microbe interaction.

## Introduction

Rhizoremediation, a technique that exploits the synergistic interactions between plants and their associated microorganisms to enhance pollutant degradation, has emerged as an attractive technology for the elimination of contaminants. Plant roots create highly active biological hotspots characterized by elevated nutrient availability, intense microbial activity and strong physicochemical gradients that stimulate biodegradation processes compared with those of bulk soils, highlighting the importance of plant–microbe interactions for remediation success (Kuiper *et al*., 2004; Glick, 2010; Bell *et al*., 2014; Handakumbura *et al*., 2021). Most current knowledge about the molecular mechanisms underlying these interactions derives from studies of plant growth-promoting rhizobacteria (PGPR), which have demonstrated the capacity of microorganisms to influence plant growth through nutrient mobilization, phytohormone production, stress alleviation and pathogen suppression (Lugtenberg and Kamilova, 2009; Backer *et al*., 2018). Increasing evidence indicates that plant-associated bacteria can profoundly reshape root development through the modulation of host signaling networks, involving hormone and signaling pathways (Vanstraelen and Benková, 2012; Kazan, 2013; Müller and Munné-Bosch, 2015). However, there is a knowledge gap about how pollutant-degrading bacteria influence plant physiology beyond their catabolic activities and this is particularly relevant because successful rhizoremediation depends not only on pollutant degradation but also on the ability of bacterial degraders to establish productive interactions with their plant hosts and surrounding microbial communities (Kuiper *et al*., 2004; Gerhardt *et al*., 2009; Molina *et al*., 2021; Mahmud *et al*., 2026).

Bacterial communication systems are likely to play a central role in the regulation of such responses. Among these, *quorum* sensing (QS) represents one of the most widespread mechanisms through which bacteria coordinate gene expression in response to environmental and population-dependent cues. In Gram-negative bacteria, QS is typically mediated by LuxI/LuxR systems where diffusible acyl-homoserine lactones (AHLs) are synthesized by LuxI proteins and detected by LuxR-family transcriptional regulators. These systems govern a wide range of functions, including biofilm formation, stress adaptation, motility, metabolism and host colonization, thereby influencing bacterial fitness across diverse environments (Papenfort and Bassler, 2016; Mukherjee and Bassler, 2019). Unlike canonical LuxR proteins, orphan-LuxR or LuxR-solo regulators lack an adjacent LuxI (Patankar and González, 2009b; Subramoni and Venturi, 2009) and are now recognized as potential environmental sensors capable of responding to signals originating from neighboring microorganisms, host organisms or other environmental sources. This exceptional versatility places orphan LuxR regulators at the interface between environmental perception and adaptive gene regulation (Venturi and Ahmer, 2015). Nevertheless, despite their widespread distribution among environmental bacteria, only a small fraction of orphan LuxR regulators have been functionally characterized, and relatively few have been implicated in regulating plant-associated phenotypes or mediating inter-kingdom communication (Ferluga and Venturi, 2009; McIntosh *et al*., 2009, 2019; Subramoni *et al*., 2011; Cellini *et al*., 2022). Whether orphan LuxR regulators from pollutant-degrading bacteria contribute to plant developmental reprogramming remains essentially unknown.

*Novosphingobium resinovorum* HR1a constitutes an excellent model for addressing these questions. This bacterium efficiently degrades a broad range of PAHs and successfully colonizes plant roots (Molina *et al*., 2021), making it a promising candidate for rhizoremediation applications. Although its catabolic capabilities have been investigated extensively (Segura *et al*., 2017), much less is known about the molecular mechanisms governing its interaction with plant hosts (Segura *et al*., 2021; Segura and Molina, 2023). In particular, it remains unclear whether bacterial regulatory networks contribute to the developmental, physiological and ecological responses observed during root colonization. LuxR874, is a previously uncharacterized orphan LuxR-family regulator from *N. resinovorum* HR1a. The amino acid sequence of LuxR874 contains signatures characteristic of Class D orphan LuxR regulators, which are found exclusively in plant-associated bacteria (PAB), where they regulate virulence-or symbiosis-related functions (Bez et al., 2023). Therefore, we hypothesized that LuxR874 regulates bacterial functions required for plant interactions during rhizoremediation.

To test this hypothesis, we combined genetic analysis, phenotypic characterization, bacterial and plant transcriptomic analysis, and physiological measurements to study the function of LuxR874 in gnotobiotic clover systems. Gnotobiotic systems provide a powerful experimental framework for dissecting specific plant–microbe interactions while maintaining biological relevance. By reducing ecological complexity of natural plant microbiomes (reviewed in Fitzpatrick et al., 2020), these systems facilitate the identification of bacterial functions that directly influence host physiology and allow the mechanistic investigation of signaling processes occurring across biological kingdoms (Vorholt *et al*., 2017; Trivedi *et al*., 2021). Our results revealed that LuxR874 inactivation modulates the expression of genes involved in aromatic and sugar metabolism. In the context of inter-kingdom signaling, LuxR874 was required for the activation of ROS-and hormone-associated host responses and for the induction of profound changes in root architecture. Together, these findings identified a previously unknown role for an orphan LuxR regulator in coordinating bacterial physiological responses with host developmental reprogramming during plant–microbe interaction.

## Experimental procedures

### Gnotobiotic system setup

Gnotobiotic systems were established as previously described (Molina *et al*., 2021). Briefly, glass jars containing 50 mL of glass beads were sterilized by double autoclaving. Each jar received 25 mL of sterile water supplemented with Fe-EDTA (125 μM) and approximately 150 mg of surface-sterilized white clover seeds (*Trifolium repens*). Seeds were surface sterilized with 70% ethanol followed by 25% sodium hypochlorite treatments as in (Molina *et al*., 2021).

*N. resinovorum* HR1a cultures were grown overnight in LB medium at 30°C with shaking (200 rpm). Cell suspensions were adjusted to approximately 10 CFU mL ¹ (OD_660_ = 0.005) in the Fe-EDTA solution prior to inoculation.

Gnotobiotic systems were incubated in a controlled growth chamber under a 16 h light period (24°C; 103–279 μmol m□² s□¹) followed by an 8 h dark period (20°C). Unless otherwise stated, plants were harvested after 8 days of incubation. Plant material was collected aseptically using sterile forceps, immediately frozen in liquid nitrogen and stored at −80°C until further processing.

For bacterial quantification, 100 μL samples from the aqueous phase of each jar were serially diluted and plated on LB agar containing rifampicin (10 μg mL ¹) to estimate inoculated HR1a populations. To estimate the number of bacteria adhered to the roots, 1 cm sections from four clover roots were washed three times with 1 mL of distilled water. Afterwards, 1 mL of sterile water and 3 glass beads were added to the tubes containing the root sections, which were then vortexed for 1 min. Serial dilutions were plated to count the colony-forming units (CFUs).

### Construction of *luxR874* mutant and complemented strains

An internal 418 bp fragment of *luxR874* (QUC32_RS13965) was amplified from genomic DNA of *N. resinovorum* HR1a using oligonucleotides 874-Fwd and 874-Rev (Suppl. Table 1). The amplified fragment was cloned into pMBL™-T, generating plasmid pMBL*874*.

The Ω-Km cassette was excised from pHP45ΩKm following digestion with *Bam*HI and *Sca*I, purified from agarose gels (Prentki and Krisch, 1984) and inserted into *Bam*HI-digested pMBL874, generating plasmid pMBL*874*Km. This construct was introduced into HR1a by electroporation. Kanamycin-resistant transformants, resulting from homologous recombination events, were selected and verified by Southern blot hybridization. The insertion within *luxR874* was additionally confirmed by sequencing bacterial isolates recovered from gnotobiotic systems inoculated with the mutant strain. The resulting mutant strain *N. resinovorum* HR1a *luxR874*::pMBL*874*Km was hereafter referred as the *luxR874* mutant.

For genetic complementation, the complete *luxR874* gene together with its native promoter region was amplified using primers Com874-PstI and Com874-EcoRI (Suppl. Table 1) and cloned into pSEVA637 (Silva-Rocha *et al*., 2013), generating plasmid p874. The resulting construct and the empty vector control (p0) were introduced by electroporation into both wild-type and *luxR874* mutant strains.

### Quantification of hydrogen peroxide accumulation

Hydrogen peroxide accumulation was determined using 3,3′-diaminobenzidine (DAB) staining.

After eight days of incubation, plants were removed from the gnotobiotic systems, weighed, and incubated overnight in a 1 mg mL ¹ DAB solution in darkness at room temperature. Following staining, chlorophyll and pigments were removed by incubation in 96% ethanol for 48 h (Molina and Kahmann, 2007; Lou *et al*., 2016).

For quantitative measurements, stained tissues were frozen in liquid nitrogen and homogenized in 0.2 M HClO□. After centrifugation at 12,000 × g for 10 min at 4°C, absorbance of the supernatants was measured at 450 nm. Hydrogen peroxide concentrations were calculated using calibration curves generated with standards ranging from 6 to 100 nmol mL□¹.

### Ethylene quantification

Ethylene production was measured using modified gnotobiotic systems established in 20 mL glass headspace vials fitted with gas-tight septa.

Inoculated and non-inoculated systems, in the presence or absence of clover plants, were incubated for 8 days under the same environmental conditions described above. Ethylene accumulation in the vial headspace was quantified using an Agilent HP4890D gas chromatograph equipped with a flame ionization detector (FID) and an HP-PLOT/Q column.

Injector, column and detector temperatures were maintained at 200°C, 40°C and 250°C, respectively. Helium was used as carrier gas at a flow rate of 5 mL min ¹ and pressure of 11.065 psi. Samples were injected automatically using a PAL RSI 85 autosampler configured for a 1.5 mL injection volume.

Ethylene concentrations were calculated using a certified 2% (v/v) ethylene standard (Air Liquide Spain). Five independent biological replicates containing approximately 20 plants each were analyzed. Following gas measurements, plants were harvested and their dry biomass was determined.

### Bacterial transcriptomic experiments

Overnight cultures grown in M9 minimal medium plus glucose of *N. resinovorum* HR1a and *luxR874* mutant were diluted in the gnotobiotic systems prepared as above to reach an initial OD_660nm_ of 0.005. The systems were placed in a growth chamber with the same conditions as above. Eight days after planting and inoculation, the liquid fractions from four gnotobiotic systems were pooled and centrifuged at 6000 rpm for 10 min. The pellets containing the cells were immediately frozen in liquid nitrogen and stored at-80°C. Two independent experimental replicates were analyzed.

For the transcriptomic experiments of the strains growing in glucose, overnight cultures were prepared as above and the next day were diluted into fresh medium (M9 minimal medium plus glucose). When the cultures reached an OD_660nm_ of 1.5, the cells were harvested by centrifugation and immediately frozen in liquid nitrogen and stored at-80°C. Three independent experimental replicates were analyzed.

Total RNA extraction and preparation of libraries were carried out as in (Molina *et al*., 2021). Briefly, the RNA was extracted using the Trizol method (TRIzol RNA Isolation Reagents, Thermo Fisher Scientific), followed by further DNAse treatment and purification with the RNeasy Mini Kit (Qiagen). One microgram of RNA per sample was used to prepare the sequencing libraries which were generated by Novogene (UK). Sequencing and downstream bioinformatics analyses were also performed by Novogene as previously described (Molina *et al*., 2021).

### Plant RNA extraction and sequencing

Total RNA was extracted from 100 mg of frozen root tissue using the NucleoBond RNA Soil Isolation Kit (Macherey-Nagel, Germany).

RNA quality and concentration were assessed using a NanoDrop™ spectrophotometer for purity, agarose gel electrophoresis for degradation check, and an Agilent 2100 Bioanalyzer (Agilent Technologies, CA, USA) to determine RNA integrity. For library preparation, polyadenylated mRNA was enriched using oligo(dT) magnetic beads, followed by fragmentation and random-primed cDNA synthesis. The final cDNA libraries underwent end-repair, A-tailing, adapter ligation, and PCR amplification. Sequencing was executed on an Illumina platform (e.g., NovaSeq 6000) using a 150 bp paired-end strategy (PE150).

### Sequence quality control, plant transcriptome reconstruction and differential expression analysis

Raw RNA sequencing reads derived from plant transcriptomes were first quality filtered and trimmed using standard preprocessing pipelines (Novogene, Germany).

Plant-derived transcripts were reconstructed from reads mapped to the *T. repens* reference genome (GCA_030408175.1; UTM_Trep_v1.0) using HISAT2 v2.2.1 (Kim *et al*., 2019). Aligned reads were sorted and processed into BAM format prior to transcript reconstruction using StringTie v2.2.1 (Kovaka *et al*., 2019). in reference-guided assembly mode. Transcript abundances were subsequently quantified with Salmon v1.10.2 (Patro *et al*., 2017) in quasi-mapping mode against the reconstructed transcriptome, generating raw count matrices for downstream analyses.

Differential expression analyses between treatments were performed using DESeq2 (Love *et al*., 2014). Statistical significance was assessed using Benjamini–Hochberg false discovery rate (FDR) correction, and genes with adjusted *p*-values < 0.05 were considered differentially expressed (Benjamini and Hochberg, 1995).

To visualize global transcriptional patterns, principal component analysis (PCA) was performed on normalized expression matrices using the *prcomp* function in R, applying unit variance scaling (scale=TRUE). The first two principal components (PC1 and PC2), which accounted for the largest proportion of variance, were retained for visualization. PCA plots were generated using *ggplot2* (Wickham, 2016) and sample labels were positioned with geom_text_repel.

### Functional enrichment analyses

We performed enrichment analyses using Pfam domains, COG functional categories, and KEGG orthology assignments derived from the clover annotation background. These complementary frameworks allowed us to summarize the broader structural, evolutionary, and biochemical signatures associated with the observed transcriptional changes, providing a higher-level view of the processes disproportionately represented among the differentially expressed genes (DEGs).

For each functional category, enrichment was assessed by comparing the number of DEGs assigned to a given category with the corresponding number of genes in the background gene set. The background comprised all annotated genes included in the transcriptomic analysis. Overrepresentation was evaluated using Fisher’s exact test based on a 2 × 2 contingency table. Enrichment magnitude was quantified using the odds ratio. To assess the predominant direction of transcriptional regulation within each enriched category, the mean log2 fold change (log_2_FC) of the DEGs contributing to each category was calculated. Categories with a positive mean log_2_FC were classified as predominantly upregulated, whereas those with a negative mean log_2_FC were classified as predominantly downregulated.

*p*-values from the enrichment analyses were corrected for multiple testing using the Benjamini– Hochberg procedure. Functional categories with an adjusted *p*-value (FDR) < 0.05 were considered significantly enriched.

## Results

### 1 A LuxR solo transcriptional regulator is required for HR1a-induced changes in root morphology

Inoculation of clover seeds with *N. resinovorum* HR1a produced a striking modification of root architecture relative to non-inoculated plants. HR1a-treated roots were shorter (Fig. 1A and D), displayed increased root diameter (Fig. 1B) and exhibited abundant root hair formation (Fig. 1C) (Molina et al., submitted). In contrast, plants inoculated with the *luxR874* mutant developed roots that closely resembled those of non-inoculated controls. Importantly, reintroduction of *luxR874* into the mutant background partially restored the wild-type phenotype, confirming that the observed developmental response was specifically associated with the function of this orphan LuxR regulator (Fig. 1A, B, C and D). These effects strictly depended on bacterial viability, as heat-killed cells failed to induce the root phenotype observed with live cultures (Fig. 1E). No significant differences in colonization levels were detected between the wild-type strain and the *luxR874* mutant (Fig. 1F).

**Figure 1:**
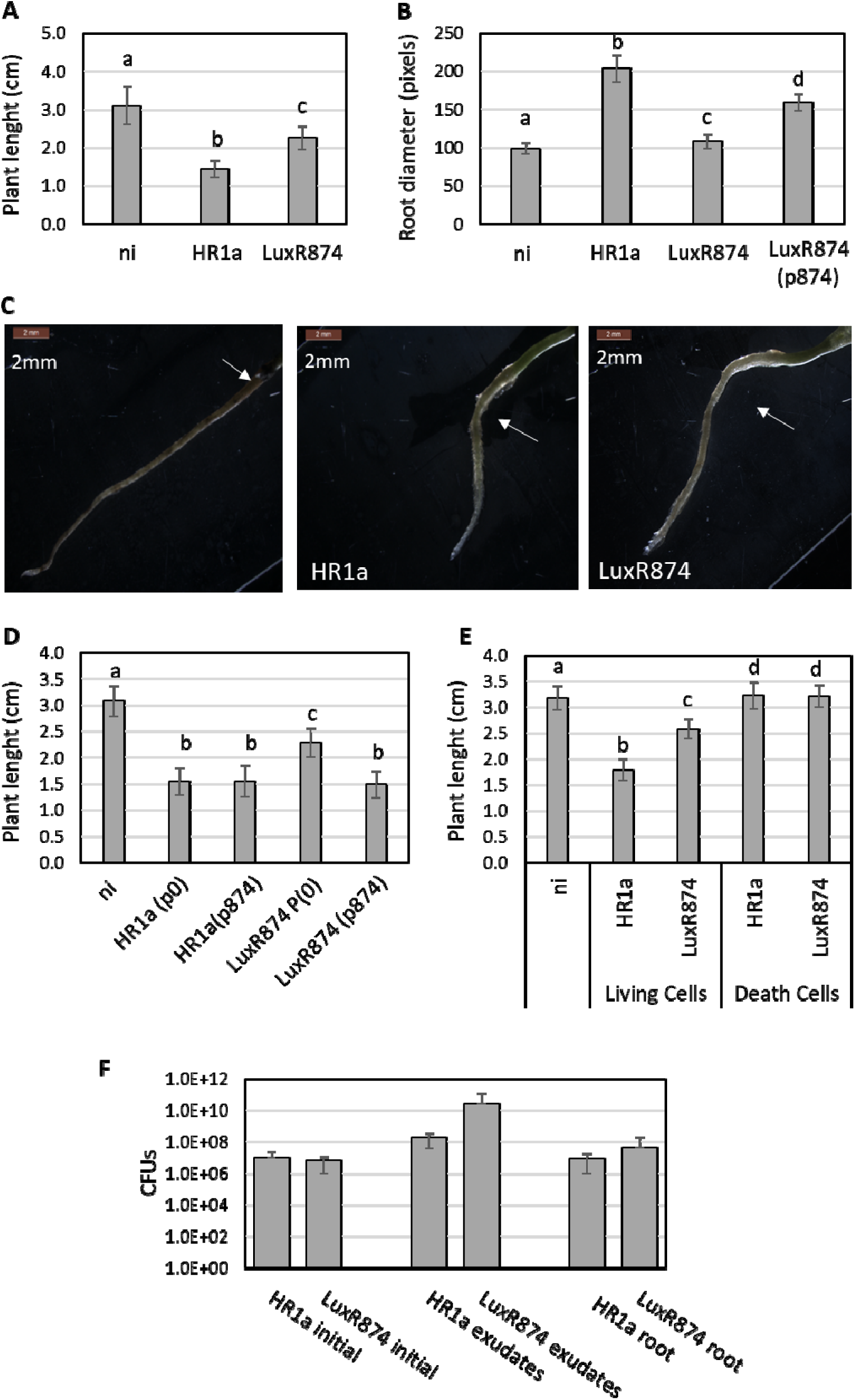
Root morphology of non-inoculated clover (ni), clover inoculated with *N. resinovorum* HR1a (HR1a) and clover inoculated with the *luxR874* mutant (LuxR874). A) Plant length after 8 days of growth in the gnotobiotic systems; (n= 60). B) Root diameter measured using ImageJ from images acquired with a Zeiss Axioskop microscope (Germany) coupled to a Nikon Digital SigH DS-L1 (Japan) (10x) (n= 12). Root diameter was measured using ImageJ software (Schindelin *et al*., 2012). C) Representative stereomicroscopic images of root morphology; white arrows indicate the interface between root and stem. D) Complementation assay showing plant length after 8 days in gnotobiotic systems inoculated with wild-type strain with the empty plasmid p0 [HR1a(p0)], with the plasmid carrying the *luxR874* gene p874 [HR1a(p874)], with the *luxR874* mutant strain (LuxR874) with p0 and with p874 (n=39). E) Plant length at 8 days in gnotobiotic systems inoculated with either with live or heat-killed bacterial cells (n > 50). F) Bacterial colonization levels of wild-type and *luxR874* mutant strains. Initial CFU mL^-1^, CFU mL^-1^ recovered from aqueous phase, and CFU cm^-1^ of roots are shown. Statistically significant differences (*p*<0.001) between groups are indicated by different letters.

These results indicate that the loss of the HR1a-induced phenotype in the mutant cannot be attributed to impaired root colonization, but rather to the absence of LuxR874-dependent functions involved in the modulation of the host responses.

### 2 *luxR874* is expressed at the onset of stationary phase and responds to quorum-sensing signals and plant exudates

When grown in minimal medium containing either 5 or 1 mM glucose as the sole carbon source, *luxR874* expression was detected at the onset of the stationary phase in both the mutant and the wild-type strains carrying the p*R874* plasmid, in which the green fluorescent protein gene (*gfp*) is expressed under the control of the *luxR874* promoter (Fig. 2A). This result suggests that LuxR874 does not autoregulate its own transcription. This observation was further confirmed by transcriptomic analysis (see below).

**Figure 2:**
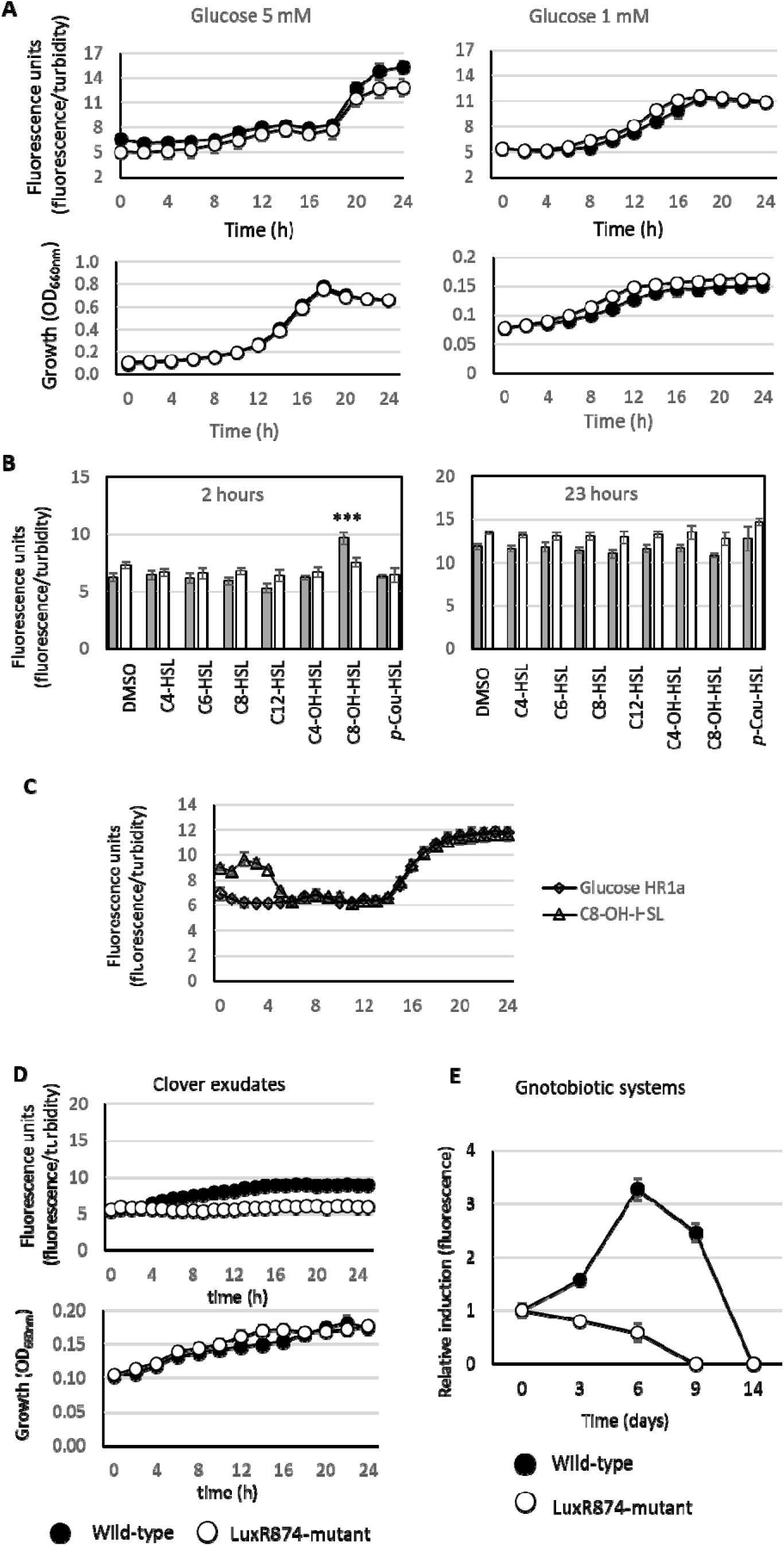
Expression of a *luxR874* promoter-*gfp* fusion under different growth conditions. A) Strain growth and relative expression (fluorescence) of the *luxR874* promoter in the wild-type (black circles) and *luxR874* mutant (white circles) strains while grown in M9 minimal medium with 5 mM or 1 mM glucose; B) Expression from the *luxR874* promoter (fluorescence) in glucose and in glucose supplemented with 10 µM of different AHLs in the wild-type (solid bars) and LuxR874 mutant (white bars) strains. Only the 2 h and 23 h time points are shown; C) Time-course of *gfp* expression from the *luxR874* promoter in the wild-type strain grown in glucose (black circles) and in glucose supplemented with 10 µM C8-OH-HSL (triangles). Fluorescence is reported as relative units normalized to optical density (OD_660nm_). *** *p*<0.001; D) GFP expression and growth of wild-type (black circles) and *luxR874* mutant (white circles) strains in clover exudates obtained 8 days after planting. E) Activity of the *luxR874* promoter during growth in gnotobiotic systems. Fluorescence is reported as relative units normalized to optical density (OD_660nm_), and bars represent mean values ± standard deviation from biological replicates.

Because expression was induced at the onset of the stationary phase, a condition often associated with quorum-sensing regulation, we investigated whether the expression of the *luxR874* gene was responsive to QS signals. To that end, we added different commercial AHLs to cultures of the wild-type strain grown with glucose as the sole carbon source. Among the tested molecules, only OH-C8-HSL induced expression from the *luxR874* promoter during the exponential growth. (Figs. 2B, C). No induction was observed in the *luxR874* mutant carrying the promoter fusion, indicating that OH-C8-HSL requires a functional LuxR874 regulator (Fig. 2B).

Next, we used clover root exudates to evaluate whether rhizosphere signals influence the expression of the *luxR874* promoter. We additionally monitored promoter activity during growth in the gnotobiotic system. In both cases, fluorescence from the *luxR874* gene was detected (Figs. 2D, 2E). In contrast, no promoter activity was observed in the *luxR874* mutant under the same conditions, suggesting that one or more rhizosphere-derived compounds may regulate *luxR874* expression.

### 3 LuxR874 coordinates aromatic compound, carbohydrate-derived and acyl-CoA-dependent metabolic pathways in *N. resinovorum* HR1a

Transcriptomic analysis comparing bacterial gene expression in gnotobiotic systems inoculated with the *luxR874* mutant with those inoculated with wild-type strain revealed that the most strongly downregulated gene clusters in the *luxR874* were associated with the metabolism and degradation of aromatic compounds and/or sugar-derivatives (Fig. 3; Suppl. Table 2).

**Figure 3:**
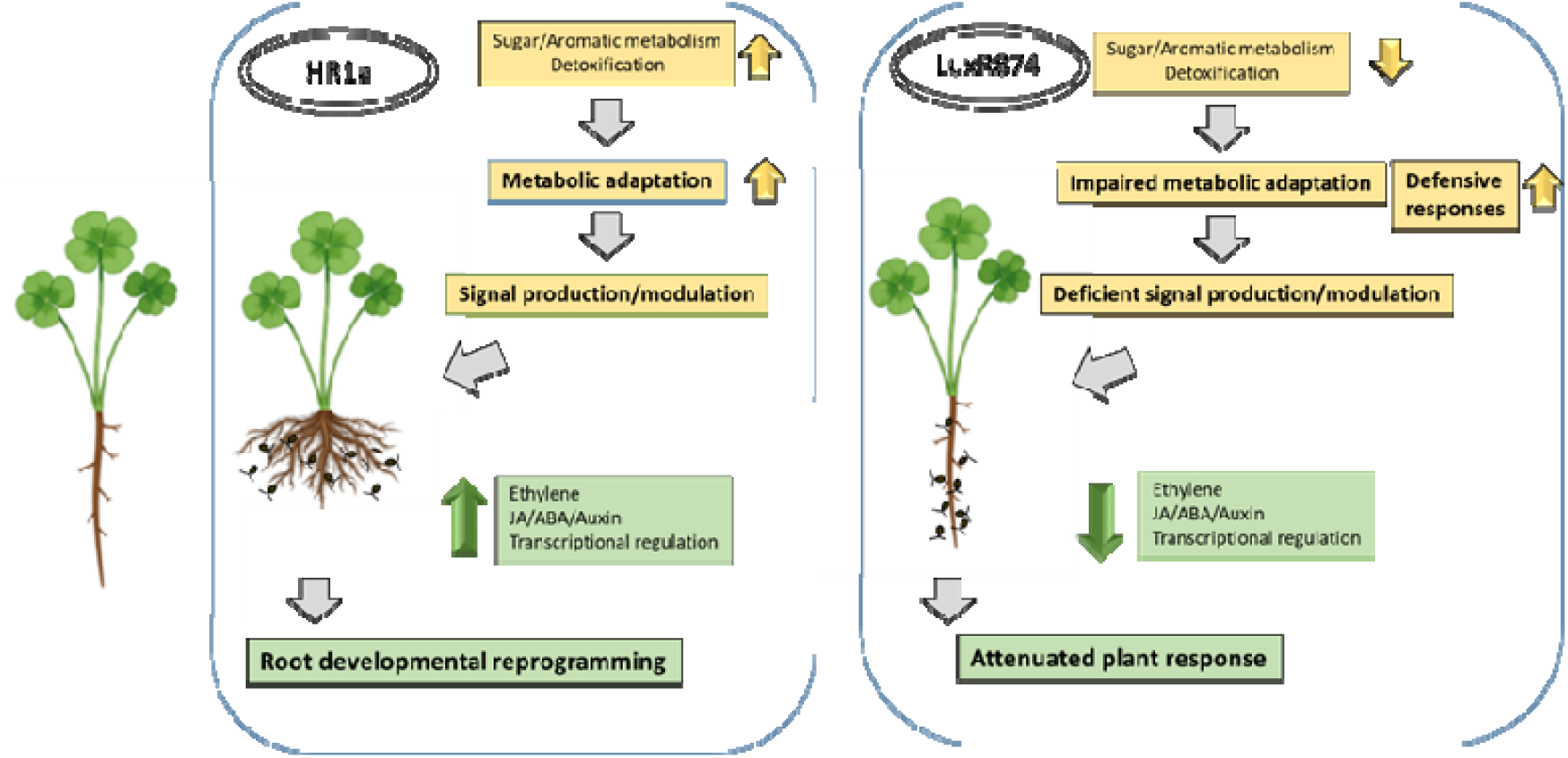
Schematic representation of the effects of the *luxR874* mutation in *N. resinovorum* HR1a and the consequences for the plant.

Genes located in close proximity to the *luxR874* gene (QUC32_RS13960 and QUC32_RS13975-QUC32_RS14015 cluster) were also expressed at higher levels in the wild-type than in the mutant strain (Suppl. Table 2). These genes encode proteins with activities related to acyl-CoA moieties, suggesting that LuxR874 influences pathways linked to fatty-acid and acyl-CoA metabolism. To validate the transcriptomic analysis results, we cloned the promoters of three downregulated genes (QUC32_RS07395, QUC32_RS06775 and QUC32_RS14740) into pSEVA637 plasmids to obtain plasmids pRS07395, pRS06775 and pRS14740 (Suppl. Table 1), thereby placing *gfp* under the control of each promoter. All three promoters were induced in the wild-type strain, whereas little or no induction was detected in the *luxR874* mutant strain, thereby corroborating the transcriptomic findings (Suppl. Fig. 1).

Because colonization by either the wild-type or *luxR874* mutant strains may differentially modify the rhizosphere environment through altered root exudation patterns, we also performed transcriptomic analysis during growth in glucose-containing medium, at the time point when *luxR874* was expressed (early stationary phase of growth). In glucose, loss of LuxR874 produced a stronger transcriptional effect in gene expression than in the gnotobiotic systems. Nevertheless, many of the highly down-regulated putative operons detected in glucose (log_2_FC ≤ - 4) were also highly down-regulated in the gnotobiotic systems (log_2_FC ≤-2.5) (Suppl. Table 2).

The pattern of gene upregulation observed in the *luxR874* mutant relative to the wild-type strain in the gnotobiotic systems was consistent with a metabolic readjustment under stress conditions. Genes involved in methionine and lipopolysaccharide biosynthesis, and transport functions, as well as three clusters of genes encoding DNA polymerase Y-related functions and other error prone polymerases, were overexpressed in the mutant strain in the presence of the plant but not during growth on glucose (Suppl. Table 3). Expression of QUC32_RS11065 was validated as described above by monitoring *gfp* expression during growth in medium containing clover root exudates (Suppl. Fig. 1).

### 4 Global transcriptional pattern of plant gene expression revealed strong differences among treatments

Principal component analysis (PCA) across treatments (non-inoculated plants, plants inoculated with the wild-type strain [HR1a], and plants inoculated with the *luxR874* mutant) revealed marked differences in transcriptome structure among treatments (Suppl. Fig. 2). Analysis of PCA loadings identified the transcripts contributing most strongly to the separation along PC1 (Suppl. Table 4), which is the axis that separates HR1a-inoculated plants from both control and *luxR874* mutant samples. This pattern suggests that the *luxR874* mutation disrupts the bacterial signals required to elicit a coordinated host transcriptional program, resulting in a transcriptomic state that resembles the heterogeneous baseline of non-inoculated plants.

Comparison between transcripts from plants inoculated with HR1a strain and non-inoculated plants (HR1a *vs* Control) revealed the most extensive transcriptional reprogramming, with 274 differentially expressed genes (DEGs), of which an overwhelming majority (256 genes) were upregulated (Suppl. Table 5). This pattern indicates a broad activation of host transcriptional programs upon colonization by the wild-type strain, consistent with HR1a acting as a potent modulator of plant physiology.

Comparison of the gene expression in plants inoculated with the *luxR874* mutant against non-inoculated plants (luxR874 *vs* Control) revealed a milder response, with 162 DEGs distributed nearly evenly between upregulated (78) and downregulated (84) genes (Suppl. Table 6). In contrast, plants inoculated with the *luxR874* mutant exhibited a substantially weaker transcriptional response when compared with clover inoculated with HR1a (luxR874 *vs* HR1a): only 90 DEGs were detected, and the expression pattern was strongly polarized: 80 were downregulated, whereas only 10 were upregulated (Suppl. Table 7). This profile indicates that the mutant retains partial ability to modulate host transcription, but lacks the strong activation signature characteristic of the WT strain.

### 5 *N. resinovorum* HR1a, but not the *luxR874* mutant, induces extensive transcriptional reprogramming in clover plants

Analysis of the DEGs in the HR1a *vs* Control contrast revealed a strong transcriptional activation in clover. Among the most strongly upregulated transcripts, we identified multiple transcription factors, several protein kinases, and genes involved in cell-wall remodeling, including pectinesterases and PMEI-like proteins. Strong induction was also observed for enzymes associated with lipid and terpene biosynthesis, such as 3-ketoacyl-CoA synthase and terpene synthase, jasmonate-related enzymes, calmodulin-binding proteins, DnaJ chaperones, and oxidoreductases (Fig 4; Suppl. Table 5).

**Figure 4.**
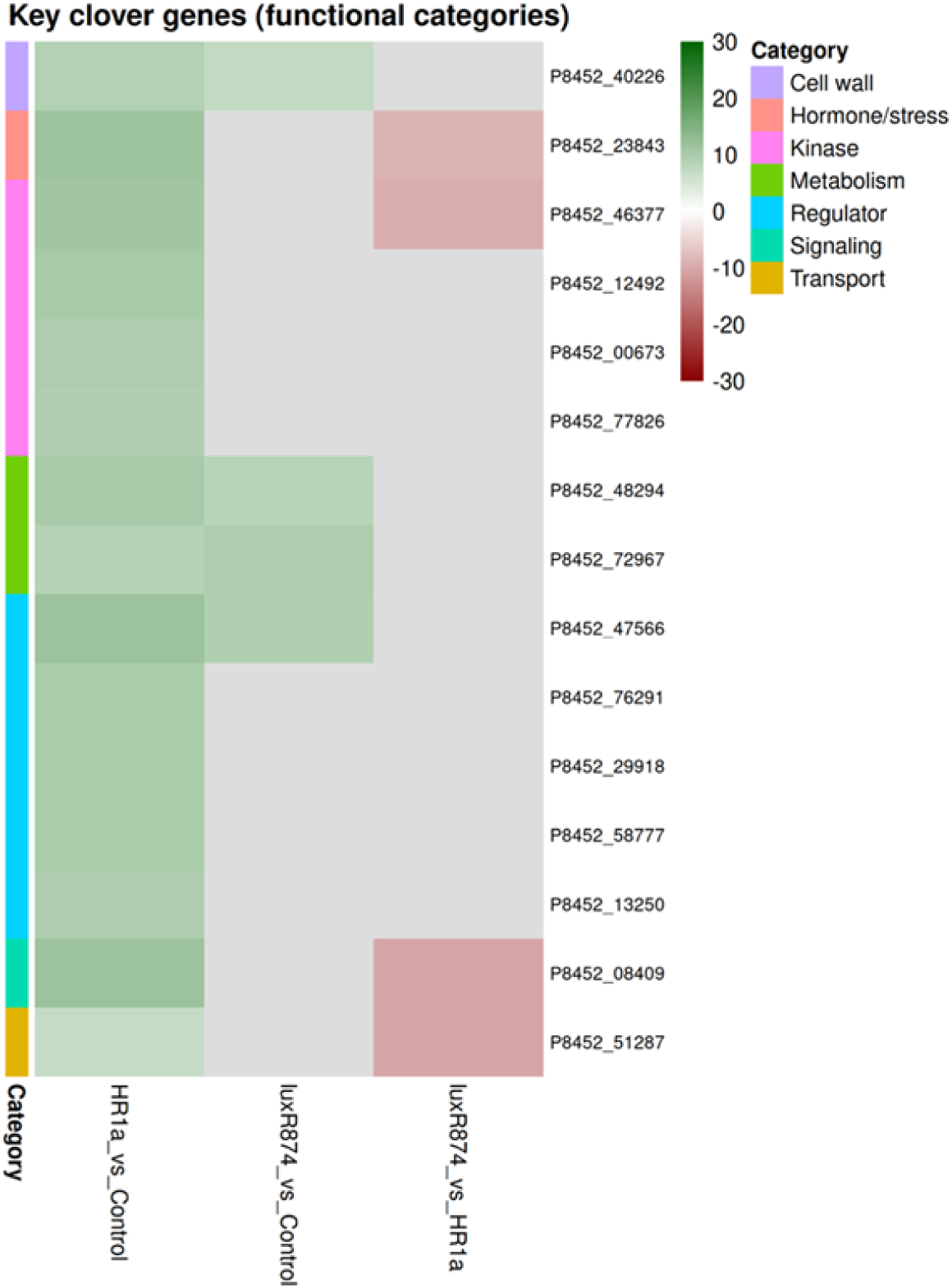
Heatmap showing the expression profiles of 15 key selected *Trifolium* genes in the contrast HR1a_*vs*_Control, and profiles of these genes in the luxR874_*vs*_Control and luxR874_*vs*_HR1a contrasts. Log□ fold-change values are displayed using a red–grey–green gradient, where red denotes transcriptional repression, green indicates induction, and grey represents near-neutral changes. Genes are grouped by functional category, including regulators, signaling components, kinases, cell-wall related genes, hormone/stress-responsive genes, transporters and specialized metabolic enzymes, and ordered accordingly to facilitate visual comparison across contrasts. Row annotations indicate the functional class of each gene, enabling rapid identification of coordinated responses within and between categories.

The luxR874 *vs* Control contrast showed a predominantly repressive transcriptional response, with many clover transcripts (84), exhibiting large negative fold-changes (Suppl. Table 6). Several of the most strongly downregulated transcripts included phosphate transporters (e.g., PT3, log□FC ≈ −23.5), sucrose synthase 6 (SUS6), 1-aminocyclopropane-1-carboxylate synthase (ACS), lipoxygenases (LOX2), and multiple LRR receptor-like kinases, indicating broad suppression of nutrient transport, carbohydrate and hormone metabolism, and cell-surface signaling components. Additional highly repressed genes included cytochrome P450 enzymes, UDP-glycosyltransferases, O-glucosyltransferases, aldo-keto reductases, and several NB-LRR resistance proteins, suggesting attenuation of diverse metabolic and defense-related pathways.

Some of the up-regulated genes in the contrast luxR874 *vs* Control were also up-regulated in the contrast HR1a *vs* Control, suggesting that there are some common plant responses to the bacterial presence, independent of the LuxR874 transcriptional regulator.

The absence of coordinated induction of cell-wall modification enzymes, hormone-related metabolic genes, and developmental regulators (e.g. Zinc-finger of the FCS-type, C2-C2, Protein TERMINAL FLOWER 1-like, NPH3 and Myb families of transcription factors), all of them highly activated when plants were inoculated with HR1a, in plants inoculated with the *luxR874* mutant (Suppl. Table 6 and 7), provides a molecular explanation for why *luxR874*-inoculated roots resemble non-inoculated controls rather than HR1a-treated plants (Fig. 1). Reduced transcription of genes involved in terpene and flavonoid biosynthesis, and disease resistance genes suggested that LuxR874-mutant does not triggered a defensive response in clover as strong as HR1a.

We examined whether these transcriptional shifts converge on specific protein domains, orthologous groups or metabolic pathways. Pfam, Gene Ontology (GO), Clusters of Orthologous Groups (COG) and KEGG enrichment analysis identified clear functional differences among treatments (Fig. 5; Suppl. Fig. 3). HR1a inoculation promoted enrichment of regulatory, signaling, and stress-related features, whereas the *luxR874* mutant elicited a markedly reduced functional response. In the luxR874 vs HR1a comparison, Pfam analysis showed photosynthesis-related domains were enriched, while multiple regulatory and stress-associated domains were underrepresented while in the other enrichment analysis no significant differences were identified (Fig. 5A and Suppl. Fig. 3).

**Figure 5.**
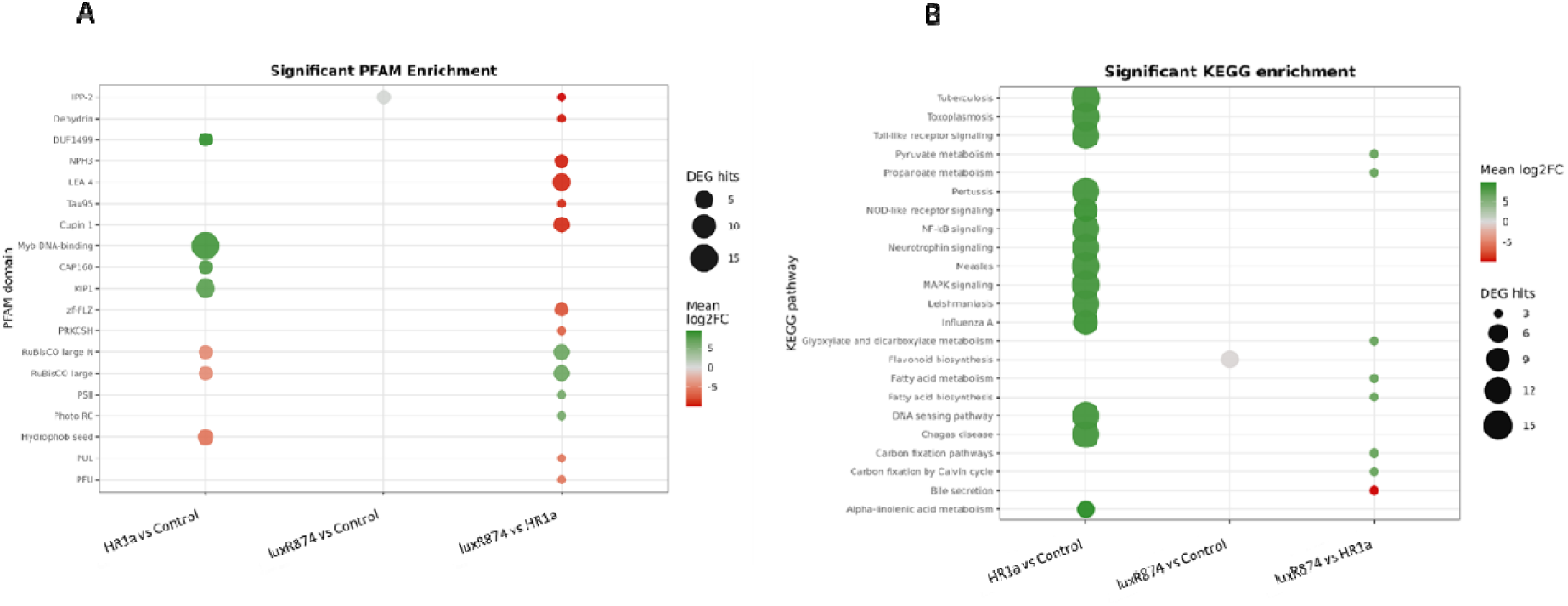
Pfam and KEGG enrichment across transcriptomic contrasts. Bubble plots summarizing Pfam protein-domain enrichment (A) and KEGG pathway enrichment (B) (FDR < 0.05) across the three transcriptomic contrasts. Bubble size represents the number of annotated differentially expressed genes, and bubble color indicates the mean log fold change. Green indicates predominantly upregulated functions, whereas red indicates predominantly downregulated functions.

These transcriptional changes highlight extensive host reprogramming triggered by HR1a, involving coordinated activation of regulatory factors, signaling components, cell-wall modification enzymes, and specialized metabolic pathways that align with the pronounced HR1a-induced root phenotype (Fig. 1). The *luxR874* mutant fails to activate these responses. This profile is consistent with its failure to induce the HR1a-specific root phenotype (Fig. 1).

### 6 Hormone-associated transcriptional reprogramming revealed by functional annotation

Focusing our attention on targeted hormone-related gene family signatures, we monitored core hormone-responsive modules, including auxin-responsive AUX/IAA and ARF domains (PF02309, PF00046), jasmonate-associated bHLH regulators (PF00010), ABA-linked LEA and dehydrin domains (PF03168), and sugar metabolism enzymes (PF00201, PF02885).

Integration of these annotations identified a comprehensive set of hormone-associated genes that were grouped into four major categories: ABA (22 genes), auxin (10 genes), jasmonate (10 genes), and sugars (5 genes). Mapping their expression profiles across contrasts revealed clear hormone-specific patterns. In HR1a_*vs.*_Control, ABA-, auxin-, and jasmonate-responsive genes showed strong activation (mean log FC > 8), consistent with extensive transcriptional reprogramming associated with stress adaptation and developmental remodeling. In contrast, luxR874_*vs.*_Control displayed marked repression of ABA-, auxin-, and sugar-associated genes, whereas luxR874_*vs.*_HR1a exhibited a broad downregulation across virtually all hormone-responsive categories, highlighting the importance of LuxR874 in activating host hormonal pathways (Fig. 6).

**Figure 6.**
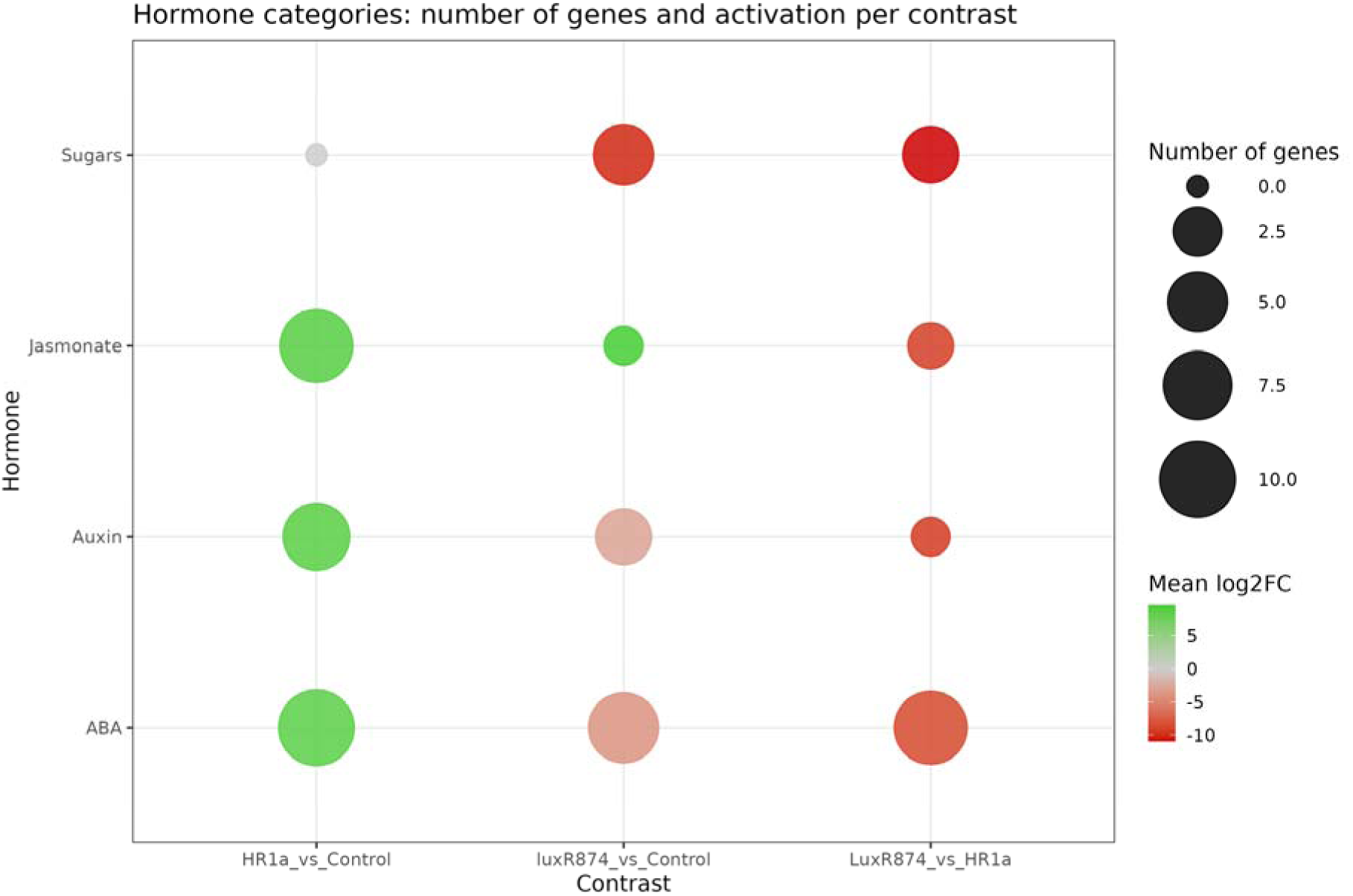
Hormonal landscape across transcriptomic contrasts. Bubble plot summarizing hormone-associated transcriptional responses identified through functional annotation (GO, KEGG and PFAM). Each bubble represents a hormone category within a specific contrast. Bubble size corresponds to the number of genes assigned to that hormone, while color reflects the mean log fold change, with green indicating activation and red indicating repression. This visualization integrates both the magnitude and direction of hormone-related transcriptional shifts, highlighting contrast-specific patterns of ABA-, auxin-, jasmonate-and sugar-responsive gene regulation.

**Figure 7:**
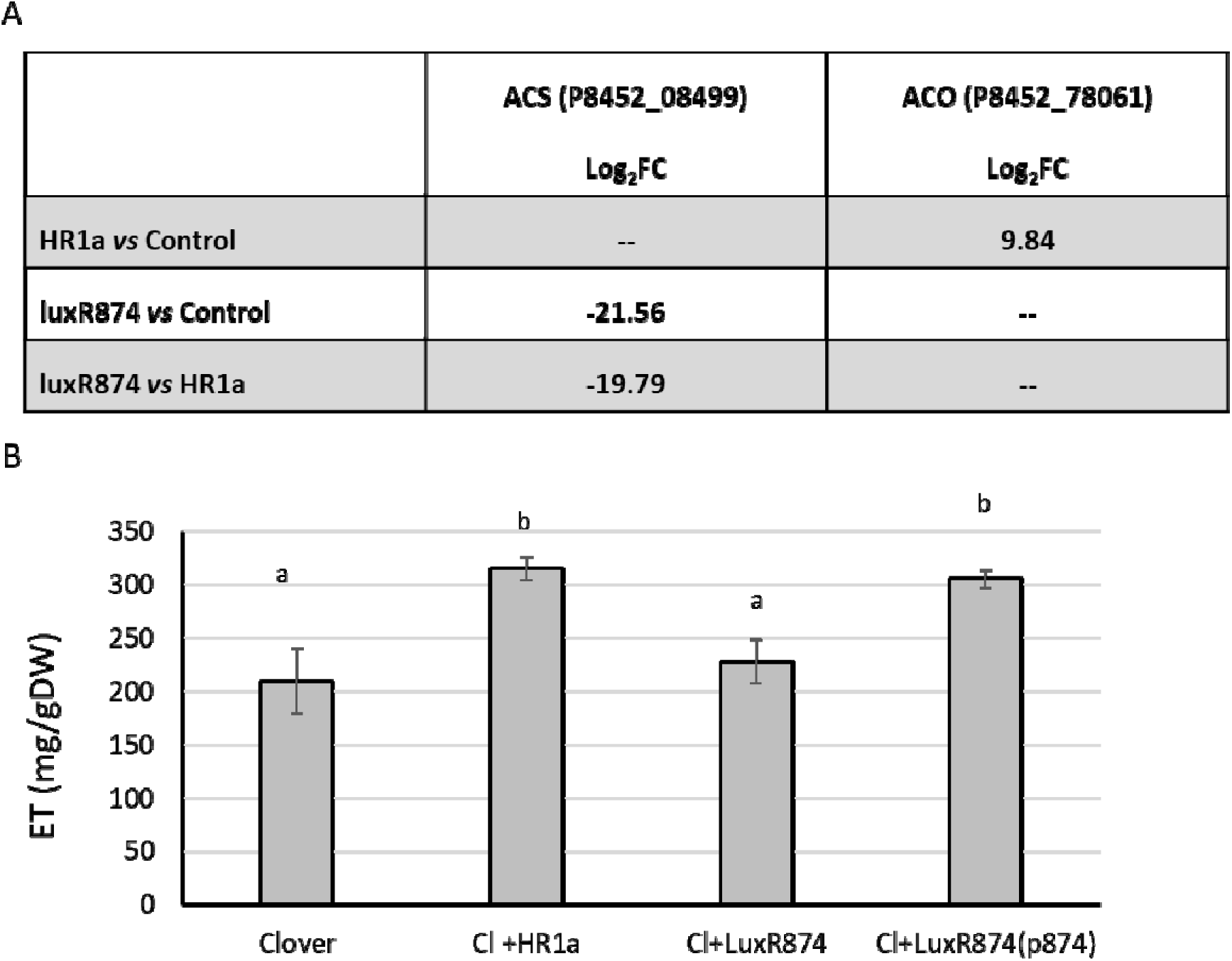
A) Differential expression of genes involved in ethylene biosynthesis. 1-aminocyclopropane-1-carboxylate synthase (ACS; P8452_08499) and ACC oxidase (ACO; P8452_78061). B) Ethylene [Et] accumulation in gnotobiotic systems after 8 days of incubation: Clover plants were grown in the absence of bacteria (Clover) or in the presence of N. resinovorum HR1a (Cl+HR1a), in the presence of the *luxR874* mutant (Cl+LuxR874) or in the presence of the complemented strain (Cl+LuxR874(p874). The different letters indicate that the concentrations are statistically significant (p< 0.005).

Beyond the global hormone categories identified by enrichment analyses, inspection of individual differentially expressed genes revealed activation of the ethylene biosynthetic pathway in HR1a-inoculated roots and repression of this pathway in plants inoculated with the *luxR874* mutant.

Transcriptomic evidence for ethylene activation by *N. resinovorum* HR1a was further supported by physiological measurements of ethylene production. Plants inoculated with the wild-type HR1a strain produced higher levels of ethylene than non-inoculated controls or plants inoculated with the *luxR874* mutant, whereas complementation with a plasmid-borne copy of *luxR874* restored ethylene production to levels comparable to those observed in the wild type. These patterns indicate that the presence of HR1a stimulates ethylene biosynthesis in a LuxR874-dependent manner and suggest that ethylene signaling represents an additional regulatory component of the host response to bacterial colonization.

### 8 LuxR874-dependent activation of ROS-associated responses in clover roots

Analysis of ROS-associated genes revealed a coordinated activation of oxidative signaling, redox regulation, oxylipin metabolism, cytochrome P450 functions, and jasmonate biosynthesis in HR1a-inoculated plants (Fig. 8A). In contrast, many of these pathways were attenuated or repressed in roots inoculated with the *luxR874* mutant. Direct comparison of *luxR874* and HR1a transcriptomes confirmed a broad suppression of ROS-and jasmonate-associated functions in the mutant, indicating that LuxR874 is required for activation of this oxidative signaling program.

**Figure 8.**
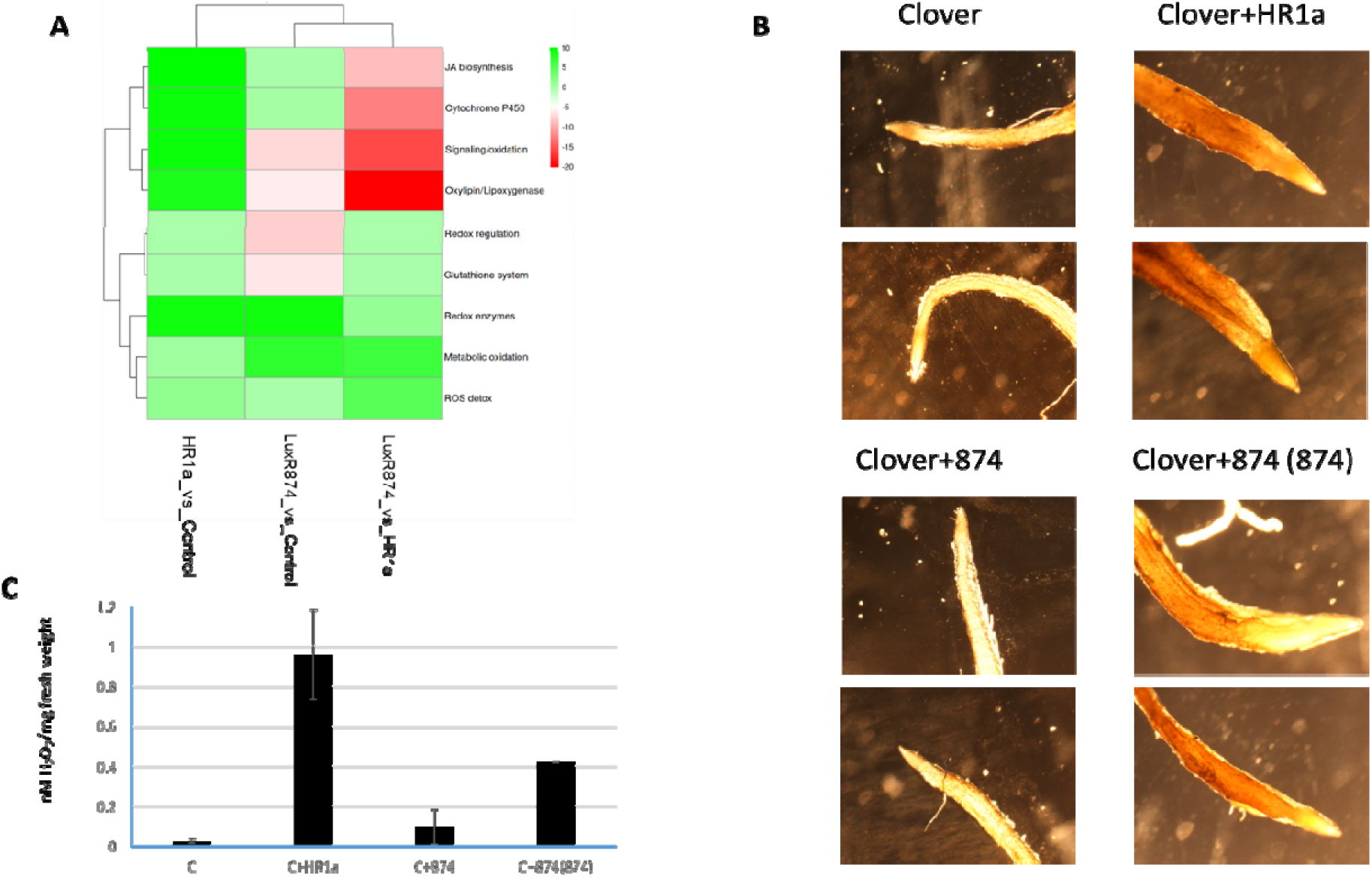
LuxR874-dependent activation of ROS-associated pathways in clover and LuxR874-dependent regulation of ROS accumulation and root morphology in clover. Heatmap showing mean log fold-change values of ROS-related functional categories across transcriptomic contrasts (A). Green indicates induction and red indicates repression. Representative images of DAB-stained roots (B) and quantification of hydrogen peroxide accumulation (C) non-inoculated plants (Clover), HR1a-inoculated plants (Clover+HR1a), plants inoculated with the *luxR874* mutant (Clover+874), and plants inoculated with the complemented strain carrying *luxR874* (Clover+874 (874)).

The ROS-associated transcriptional signatures were validated at the physiological level through DAB staining and complementation experiments. Wild-type HR1a induced strong H O accumulation in clover roots, whereas the *luxR874* mutant exhibited substantially reduced ROS levels. Importantly, reintroduction of the *luxR874* gene restored H O accumulation, demonstrating that the observed phenotype is directly dependent on this regulator. These findings provide functional evidence that LuxR874 regulates bacterial functions required to trigger oxidative signaling in the host and suggest that ROS production represents a key component of the transcriptional and morphological reprogramming induced by HR1a.

## Discussion

Orphan LuxR regulators have emerged as remarkably diverse transcriptional regulators capable of responding to bacterial, plant-derived and other, yet unidentified, environmental signals (Patankar and González, 2009b). Although members of the PAB LuxR-solo subfamily have been implicated in bacterial virulence, stress adaptation, rhizosphere competence and host colonization (Patankar and González, 2009a; Subramoni *et al*., 2011), the functional characterization of orphan LuxR regulators has been largely focused on bacterial phenotypes, including motility, biofilm formation, stress tolerance and colonization capacity (Zhang *et al*., 2007; Subramoni *et al*., 2011; Schaefer *et al*., 2016; Coutinho *et al*., 2018; Reina *et al*., 2019; Mosquito *et al*., 2020). Their contribution to host physiological responses remains much less understood. Our study extends this perspective by showing that an orphan LuxR regulator, LuxR874, influences the ability of *Novosphingobium resinovorum* HR1a, to induce extensive developmental and physiological responses in its host, the clover plant, revealing a broader role for this family of regulators in interkingdom communication than previously appreciated.

The bacterial transcriptomic data indicate that LuxR874 is a positive regulator of genes associated with aromatic-and sugar-derived compounds metabolism, and acyl-CoA-associated pathways. In the rhizosphere, loss of LuxR874 was additionally associated with increased expression of functions related to stress-tolerance (Fig. 3). This data suggests that LuxR874 contributes to the physiological adaptation of *N. resinovorum* HR1a across different environmental conditions (laboratory or rhizosphere).

Although both bacteria were able to colonize clover roots, HR1a induced a pronounced developmental phenotype characterized by shorter roots and increased root hair formation, whereas plants inoculated with the *luxR874* mutant remained much similar to the control phenotype. This distinction was reflected at the transcriptomic level. *N. resinovorum* HR1a induced extensive changes in regulatory, signaling, defense-associated and hormonal functions. The differences observed between wild-type and mutant strains are consistent with the contrasting plant phenotypes and indicate that bacterial colonization alone is insufficient to explain the developmental response. Rather, LuxR874-dependent bacterial functions appear to determine whether root colonization is accompanied by extensive host transcriptional reprogramming.

One possible explanation involves the interaction between LuxR874-dependent bacterial physiology and chemical signal/s present in the rhizosphere. The regulation of fatty acid degradation and lactonase-associated functions by LuxR874 is noteworthy because signaling molecules related to HSLs have been shown to influence root architecture. Medium chain HSLs reduce primary root growth while promoting lateral root and root hair formation, as well as activating ROS-, kinase-, transcription factor-and hormone-associated pathways (Ortíz Castro *et al*., 2008; Bai *et al*., 2012; Miao *et al*., 2012; Schenk *et al*., 2014; Zhao *et al*., 2016; Du *et al*., 2021; Liu *et al*., 2022). These responses closely resemble those induced in clover by the HR1a strain. In our system, C8-OH-HSL induced the early expression of the *luxR87*4 in a LuxR874-dependent mode. LuxR874 was also required for its own induction in the rhizosphere, but not under laboratory conditions, suggesting the presence of rhizospheric-derived signals that regulate *luxR874* expression. Since PAB LuxR-solo can respond to structurally diverse compounds (Schaefer *et al*., 2008; Subramoni *et al*., 2011; Mull *et al*., 2018), LuxR874 may contribute to the integration of environmental sensing, bacterial physiological state and plant development responses.

The plant transcriptomic data further support this interpretation. Our results indicate that HR1a activates jasmonate-, ethylene-, and ROS-associated pathways, whereas disruption of *luxR874* markedly attenuated these responses and reduced H_2_O_2_ accumulation, with concomitant loss of the characteristic HR1a-induced root phenotype (Fig. 3). These signaling networks are intensively interconnected and regulate of root development, cell expansion and differentiation, and environmental stress adaptation (Pitts *et al*., 1998; Singh *et al*., 2016; Qin *et al*., 2019; Bali, 2025; Sun *et al*., 2026). The observed activation of ROS and ethylene-responses in HR1a-inoculated plants provides physiological support for the transcriptomic predictions and suggest that LuxR874-dependent bacterial functions are required to elicit this coordinated plant response.

Taken together, the bacterial and plant datasets support a model in which LuxR874 contributes to a rhizosphere-associated physiological state that enables host developmental reprogramming. LuxR874 may itself respond to one or more compounds present in the rhizosphere, including plant-derived molecules, and thereby alter bacterial metabolism and physiology. Nevertheless, our results place LuxR874 at the interface between environmental sensing and host modulation and show that bacterial colonization can be uncoupled from the capacity to induce developmental responses. This finding highlights the importance of bacterial regulatory networks in shaping plant phenotypes during plant-microbe interactions.

Root modifications induced by beneficial rhizobacteria and symbiotic microorganisms have been widely reported, these microorganisms mainly provoked increases in the root surface, creating new spaces for nutrient exchange and microbial colonization (Concha & Doerner, 2020; Roca et al., 2013; Verbon & Liberman, 2016). Our results suggest that environmental degraders, such as *N. resinovorum* HR1a, and not only PGPRs, can also influence host development programs, through regulatory processes that are not necessarily linked directly to its catabolic activity. Thus, the ecological function of contaminant-degrading bacteria may extend beyond pollutant removal to include regulation of the plant phenotype in ways that could influence the establishment and functioning of the plant–microbe partnership.

## Conclusions

Our results identify LuxR874 as a key regulator linking bacterial physiological adaptation with host developmental responses during a rhizosphere interaction. Although disruption of *luxR874* did not affect root colonization, it strongly impaired the ability of *N. resinovorum* HR1a to induce root architectural changes, ROS accumulation, ethylene-related responses, and the extensive transcriptional reprogramming observed in clover plants inoculated with the wild-type strain. These findings demonstrate that successful root colonization can be uncoupled from the capacity to modulate host physiology and place LuxR874 at the interface between environmental sensing and plant response. The transcriptional signatures observed in both partners further suggest that LuxR874 contributes to the establishment of a rhizosphere-associated bacterial state that promotes host developmental reprogramming, potentially through the perception and integration of rhizospheric signals. More broadly, our study expands the functional significance of orphan LuxR regulators beyond bacterial adaptation and highlights that pollutant-degrading bacteria can influence plant developmental programs through specific regulatory networks. These findings underscore the importance of bacterial regulatory mechanisms in shaping plant-microbe interactions and suggest that the ecological contribution of environmental degraders may extend beyond pollutant removal to include modulation of host phenotype and rhizosphere functioning.

## Author Contributions

L.M. participated in the conceptualization, methodology design, research, analysis, data curation, and writing of original draft, and review and editing of final manuscript. I.H. contributed with technical work. Z. U. participated in the formal analysis, software, data curation, manuscript review and editing. A. S. participated in the conceptualization, funding acquisition, supervision, project administration, discussion of results and writing, review and editing.

All authors reviewed the manuscript, and approved the final version for publication.

## Conflicts of interest

None declared.

## Funding

This work was supported by grant number BIO2017-85994-P financed by MCIN/AEI/10.13039/501100011033/ and by “ERDF A way of making Europe” and Grant PID2020-116766GB-I00 and PID2023-148648NB-I00 financed by MCIN/ AEI/10.13039/501100011033.

## Data availability

Raw white clover RNA sequencing reads have been deposited in the NCBI Sequence Read Archive (SRA) under BioProject accession PRJNA1513061. Bacterial RNA sequencing reads have been deposited in the NCBI Gene Expression Omnibus (GEO) under accession GSE350117 and will be made publicly available upon publication of this article.

## Notes

### Competing Interest Statement

The authors have declared no competing interest.

