## Supplementary material for "An orphan LuxR regulator controls plant developmental reprogramming during interaction with *Novosphingobium resinovorum* HR1a": Supp Table 1

**Suppl. Table 1**: List of strains, plasmids and primers used in this study.

| ***Strains*** | ***Relevant proteperties*** | ***Reference*** |
| --- | --- | --- |
| *Novosphingobium* *resinovorum* HR1a | PAH degrader | Segura et al., 2018 |
| *N. resinovorum* HR1a  *luxR874*::pMBL874Km | Mutant in the *luxR874* gene | Molina et al., 2024 |
| *N. resinovorum* HR1a (pR874) | Reporter strain (wild-type) with the *luxR874* promoter driving the expression of the GFP protein | Molina et al., 2024 |
| *N. resinovorum* HR1a *luxR874::pMBL874Km mutant* (pR874) | Reporter strain (LuxR874 mutant) with the *luxR874* promoter driving the expression of the GFP protein | Molina et al., 2024 |
| ***Plasmids*** |  | ***Relevant properties*** |
| pSEVA637 | Gm^R^, promoterless *gfp* | Silva-Rocha et al., 2013 |
| p874 | pSEVA637 with the *luxR874* geneand its own promoter, used in complementation experiments | This study |
| p*R874* | pSEVA637 with the *luxR874* promoter driving the *gfp* expression | Molina et al., 2024 |
| pRS07395 | pSEVA637 with the promoter  of the gene QUC32_RS07395 driving the *gfp* expression | This study |
| pRS06775 | pSEVA637 with the promoter  of the gene QUC32_RS06775 driving the *gfp* expression | This study |
| pRS14740 | pSEVA637 with the promoter  of the gene QUC32_RS14740 driving the *gfp* expression | This study |
| pRS11065 | pSEVA637 with the promoter  of the gene QUC32_RS11065 driving the *gfp* expression | This study |
| ***Primers*** |  | ***Sequence (5´-3´)*** |
| 874-Fwd | For amplification of a *luxR874* internal fragment for mutant construction | GACATTGCCGACAAGCGTA |
| 874-Rev | For amplification of a *luxR874* internal fragment for mutant construction | CCATGACCTGCACATACGTC |
| Com874-*Pst*I | For amplification of *luxR874* with its own promoter | *ctgcag*catttcctcgtcttcg |
| Com874-*EcoR*I | For amplification of *luxR874* with its own promoter | *GAATTC*AGTATCCAGGAG |
| 874proEco | For *luxR874* promoter amplification | *GGATTC*AGTCCACTCCCGCAGGGATG |
| 874proPst | For *luxR874* promoter amplification | AA*CTGCAG*TGATGTCCTCTCGAGGCT |
| EcoR1 Pr4806 | For amplification of the QUC32_RS07395 promoter | *GAATTC*GCCGTCTCCCAGTCGCC |
| Pst1 Pr4806 | For amplification of the QUC32_RS07395 promoter | *CTGCAG*CCCCCTTGCTTACCCGC |
| EcoR1 Pr4931 | For amplification of the QUC32_RS06775 promoter | *GAATTC*GTAGCTTTCCGTATCGTC |
| Pst1 Pr4931 | For amplification of the QUC32_RS06775 promoter | *CTGCAG*CAGTTTATCCTCCCGGTG |
| EcoR1 Pr1028 | For amplification of the QUC32_RS14740 promoter | *GAATTC*CATCGTCCTTCCCCCGTATC |
| Pst1 Pr1028 | For amplification of the QUC32_RS14740 promoter | *CTGCAG*GGACAGATTGCTCGCGTGC |
| EcoR1 Pr294 | For amplification of the QUC32_RS11065 promoter | *GAATTC*GAAGGTCACCGCGACGAT |
| Pst1 Pr294 | For amplification of the QUC32_RS11065 promoter | *CTGCAG*GGATCCGTCCCCCGTTGG |
