## Supplementary material for "An orphan LuxR regulator controls plant developmental reprogramming during interaction with *Novosphingobium resinovorum* HR1a": Supp Table 2

**Suppl. Table 2**: **List of genes down-regulated (log_2_FC ≤ -2 in gnotobiotic systems and log_2_FC ≤ -4 in glucose) in the mutant strain when compared with the wild-type**. Genes downregulated in both conditions and genes contiguous to those down-regulated have been included. Chr refers to the chromosome number and pHR1a to a plasmid.

| **Gene ID** | **log_2_FC>2**  **gnot** | **log_2_FC >4**  **gluc** | **Strand** | **Gene_description** | **Chr** |
| --- | --- | --- | --- | --- | --- |
| QUC32_RS07405 | **-3.5** | -3.8 | + | gluconolactonase | 1 |
| QUC32_RS07400 | **-5.8** | **-5.4** | + | PF12680:SnoaL-like domain |  |
| QUC32_RS07395 | **-4.1** | **-6.1** | - | NADP-dependent oxidoreductase:Zinc-binding dehydrogenase |  |
| QUC32_RS07390 | **-2.1** | -1.8 | - | Rieske 2Fe-2S domain-containing protein |  |
| QUC32_RS07385 | **-4.4** | **-5.1** | - | SnoaL-like domain |  |
| QUC32_RS06775 | **-4.0** | **-4.2** | - | TonB-dependent receptor | 1 |
| QUC32_RS06770 | **-4.4** | **-4.3** | - | Glucose 1-dehydrogenase |  |
| QUC32_RS06760 | -1.6 | **-4.1** | + | ThuA domain-containing protein:Trehalose utilisation |  |
| QUC32_RS06755 | **-2.7** | **-5.3** | + | NADP-dependent oxidoreductase:Zinc-binding dehydrogenase |  |
| QUC32_RS06750 | **-3.6** | -2.3 | - | TonB-dependent receptor |  |
| QUC32_RS14725 | **--** | -2.1 | - | SnoaL-like domain | 2 |
| QUC32_RS14730 | **--** | -2.1 | - | SDR family NAD(P)-dependent oxidoreductase |  |
| QUC32_RS14740 | **-3.3** | **-5.3** | + | PF13577:SnoaL-like domain | 2 |
| QUC32_RS14745 | **-4.4** | **-6.0** | + | Aromatic ring-hydroxylating dioxygenase subunit alpha domain |  |
| QUC32_RS14750 | **-3.7** | **-6.5** | + | PF13577:SnoaL-like domain |  |
| QUC32_RS14755 | **-3.2** | -3.0 | - | Antibiotic biosynthesis monooxygenase |  |
| QUC32_RS14760 | **-2.5** |  | + | Glucose 1-dehydrogenase |  |
| QUC32_RS19000 | **-3.3** | **-5.5** | - | Antibiotic biosynthesis monooxygenase | 2 |
| QUC32_RS19005 | **-4.2** | -3.8 | - | Dabb family protein |  |
| QUC32_RS03700 | **-4.1** | **-5.3** |  | Aromatic ring-hydroxylating dioxygenase subunit alpha | 1 |
| QUC32_RS03705 | **-3.3** | **-5.1** |  | SDR family oxidoreductase |  |
| QUC32_RS06410 | **--** | -3.9 |  | EthD domain-containing protein | 1 |
| QUC32_RS06405 | **--** | **-4.1** | + | sugar phosphate isomerase |  |
| QUC32_RS06400 | **-3.8** | **-4.6** | + | Ring hydroxylating alpha subunit (catalytic domain)\| |  |
| QUC32_RS06395 |  | -3.3 | - | Medium-chain fatty-acid--CoA ligase |  |
| QUC32_RS06390 | **-2.0** |  | + | SDR family oxidoreductase |  |
| QUC32_RS06385 | -1.9 | -2.9 | - | SDR family oxidoreductase |  |
| QUC32_RS18225 | **-3.6** | **-6.3** | + | FAD-binding protein | 2 |
| QUC32_RS13960 | **-3.2** |  | - | Acyl-CoA dehydrogenase family protein | 2 |
| QUC32_RS13975 | **-3.3** | **-4.7** | + | OB-fold domain-containing protein, acyl-CoA-associated | 2 |
| QUC32_RS13980 | **-2.5** | -3.8 | + | Thiolase family protein |  |
| QUC32_RS13985 | -1.6 | -3.5 | + | Acyl-CoA dehydrogenase family protein |  |
| QUC32_RS13990 | -1.6 | **-4.1** | + | Thiamine pyrophosphate-dependent dehydrogenase E1 component subunit alpha |  |
| QUC32_RS13995 | -1.7 | **-4.1** | + | Alpha-ketoacid dehydrogenase subunit beta |  |
| QUC32_RS14000 | -1.9 | -3.8 | + | 2-oxo acid dehydrogenase subunit E2 |  |
| QUC32_RS14005 | -1.5 | -3.8 | + | VOC family protein |  |
| QUC32_RS14010 | -- | -3.7 | + | Acyl-CoA dehydrogenase |  |
| QUC32_RS14015 | -- | -3.6 | + | Phosphotransferase/ Acyl-CoA dehydrogenase |  |
| QUC32_RS27995 | **-3.2** | **-5.6** | + | Amidohydrolase family protein | pHR1a |
| QUC32_RS02575 |  | -1.5 | - | FAD-binding protein | 1 |
| QUC32_RS02580 |  | **-2.1** | - | SnoaL-like domain |  |
| QUC32_RS02585 |  | -1.6 | - | SDR family oxidoreductase |  |
| QUC32_RS02590 | **-2.4** | **-2.6** | - | Aromatic ring-hydroxylating dioxygenase subunit |  |
| QUC32_RS02595 |  | **3.3** | - | SnoaL-like domain |  |
| QUC32_RS02600 | **-2.8** | **-3.8** | - | TonB-dependent receptor |  |
| QUC32_RS05540 | **-2.2** | **-4.8** | + | ThuA domain-containing protein && PF06283:Trehalose utilisation | 1 |
| QUC32_RS05535 | **-2.6** | **-4.9** | + | Aromatic ring-hydroxylating dioxygenase subunit alpha |  |
| QUC32_RS05530 | **-2.5** | **-2.2** | - | SDR family oxidoreductase |  |
| QUC32_RS05525 | **-2.0** | **-2.8** | - | Glucose 1-dehydrogenase |  |
| QUC32_RS05520 | -1.7 | **-2.5** | - | SDR family oxidoreductase |  |
| QUC32_RS05515 | -1.8 | **--2.8** | - | VOC family protein |  |
| QUC32_RS04560 | -1.8 | -- | + | potassium-transporting ATPase subunit KdpA | 1 |
| QUC32_RS04565 | **-2.2** | -- | + | potassium-transporting ATPase subunit KdpB |  |
| QUC32_RS07270 | **-2.1** | -- | + | glucose/quinate/shikimate family membrane-bound PQQ-dependent dehydrogenase | 1 |
| QUC32_RS07265 | -1.5 | -- | - | NAD(P)/FAD-dependent oxidoreductase |  |
| QUC32_RS11980 | -1.8 | -- | - | hypothetical protein | 2 |
| QUC32_RS11980 | **-2.0** | -- | - | toxin-antitoxin system HicB family antitoxin |  |
| QUC32_RS11980 | -1.7 | -- | - | SPFH domain-containing protein |  |
| QUC32_RS17820 | **-2.0** | -- |  | helix-turn-helix transcriptional regulator | 2 |
| QUC32_RS12885 | **--** | **-4.6** | - | hypothetical protein | 2 |
| QUC32_RS22765 | **--** | **-4.3** | - | GTP cyclohydrolase II | 2 |
| QUC32_RS22770 | **--** | -2.9 | - | exodeoxyribonuclease III |  |
| QUC32_RS01905 | **--** | -2.2 | - | LLM class flavin-dependent oxidoreductase, alkanesulfonate monooxygenase | 1 |
| QUC32_RS01910 | **--** | -3.7 | - | LLM class flavin-dependent oxidoreductase, alkanesulfonate monooxygenase |  |
| QUC32_RS01925 | **--** | **-4.2** | + | PF13347:MFS/sugar transport protein | 1 |
| QUC32_RS01935 | **--** | -3.4 | - | alpha/beta hydrolase |  |
| QUC32_RS10790 | **--** | -4.1 |  | HPP family protein Transmembrane protein | 2 |
| QUC32_RS25470 | **--** | -2.0 | - | FixH family protein | 2 |
| QUC32_RS25475 | **--** | -3.2 | - | cytochrome c oxidase accessory protein FixG |  |
| QUC32_RS25480 | **--** | -3.7 | - | cytochrome-c oxidase cbb3-type subunit III FixP |  |
| QUC32_RS25485 | **--** | -3.5 | - | cbb3-type cytochrome c oxidase subunit 3 FixQ |  |
| QUC32_RS25490 | **--** | -3.8 | - | cbb3-type cytochrome c oxidase subunit 3 FixO |  |
| QUC32_RS25495 | **--** | **-4.1** | - | cytochrome-c oxidase cbb3-type subunit FixN |  |
| QUC32_RS25500 | **--** | -2.8 | + | OmpW family outer membrane protein |  |
