## Supplementary material for "An orphan LuxR regulator controls plant developmental reprogramming during interaction with *Novosphingobium resinovorum* HR1a": Supp Table 3

**Suppl. Table 3: List of genes up-regulated (log_2_foldchange (log2FC ≥ 2) in the mutant.** Some genes, contiguous to those up-regulated, have been included. Other genes with log_2_FC lower than 1.5 have been included because of the redundancy of expression of the cassette error-prone DNA polymerase/DNA polymerase Y/ImuA. Chr refers to the chromosome number and pHR1a to a plasmid.

| **GenBank ID** | **log_2_FC**  **gnot** | **Log_2_FC**  **gluc** | **Strand** | **Gene_description** | **Chr** |
| --- | --- | --- | --- | --- | --- |
| QUC32_RS23785 | **6.0** | 1.8 | + | Homocysteine S-methyltransferase family protein | 2 |
| QUC32_RS23790 | **4.9** | 1.57 | + | GNAT family N-acetyltransferase |  |
| QUC32_RS23795 | **4.0** |  | + | Vitamin B12 dependent methionine synthase, activation domain |  |
| QUC32_RS11065 | **2.7** |  | + | Porin | 2 |
| QUC32_RS11070 | 1.9 |  | + | Cytochrome b |  |
| QUC32_RS11075 | 1.6 |  | + | TonB-dependent receptor |  |
| QUC32_RS19830 | 1.8 |  | + | RES family NAD+ phosphorylase | 2 |
| QUC32_RS19835 | **2.7** |  | - | DUF2384 domain-containing protein; MbcA/ParS/Xre antitoxin |  |
| QUC32_RS00755 | **2.5** |  | - | Hypothetical protein | 1 |
| QUC32_RS03475 | 1.5 |  | - | Hypothetical protein | 1 |
| QUC32_RS03480 | 1.2 |  | - | Error-prone DNA polymerase |  |
| QUC32_RS03485 | **2.1** |  | - | DNA polymerase Y family protein |  |
| QUC32_RS03490 | **2.4** |  | - | Protein ImuA |  |
| QUC32_RS02345 | 1.6 |  | - | error-prone DNA polymerase :Bacterial DNA polymerase III alpha subunit | 1 |
| QUC32_RS02350 | **2.4** |  | - | DNA polymerase Y family protein:impB/mucB/samB family |  |
| QUC32_RS02355 | 1.8 |  | - | protein ImuA |  |
| QUC32_RS09460 | **2.3** |  | + | Hypothetical protein | 1 |
| QUC32_RS08960 | 2.2 |  | + | Cation efflux family | 1 |
| QUC32_RS10065 | **2.1** |  | - | Hypothetical protein | 2 |
| QUC32_RS10070 | 1.8 |  | - | CpsD/CapB family tyrosine-protein kinase |  |
| QUC32_RS10075 | 1.7 |  | - | Lipopolysaccharide biosynthesis protein:Chain length determinant protein |  |
| QUC32_RS10085 | **2.2** |  | - | Polysaccharide biosynthesis/export family protein |  |
| QUC32_RS11120 | **2.1** |  | + | Hypothetical protein | 2 |
| QUC32_RS16070 | 1.6 |  | + | Transcriptional regulator PadR-like family | 2 |
| QUC32_RS16075 | **2.0** |  | + | DUF2218 domain-containing protein |  |
| QUC32_RS03855 | **2.0** |  | + | RNA polymerase sigma factor (RpoE) | 1 |
| QUC32_RS29845 | **2.0** |  | - | error-prone DNA polymerase :Bacterial DNA polymerase III alpha subunit | pHR1a |
| QUC32_RS29850 | 1.8 |  | - | DNA polymerase Y family protein:impB/mucB/samB family |  |
| QUC32_RS29855 | 1.3 |  | - | Protein ImuA |  |
| QUC32_RS01025 |  | 1.86 | - | hypothetical protein- | 1 |
| QUC32_RS01030 |  | **2.38** | - | glycosyltransferase |  |
| QUC32_RS01035 |  | **2.50** | - | DUF2141 domain containing protein |  |
| QUC32_RS01040 |  | **3.03** | - | hypothetical protein |  |
| QUC32_RS01045 |  | **3.23** | - | glycosyltransferase |  |
| QUC32_RS01070 |  | 1.88 | - | hypothetical protein | 1 |
| QUC32_RS01075 |  | **2.63** | + | endonuclease/exonuclease/phosphatase family |  |
| QUC32_RS06000 |  | **2.14** | - | carbohydrate porin | 1 |
| QUC32_RS10915 |  | **2.13** | - | VOC family protein | 2 |
| QUC32_RS11710 |  | 1.54 |  | translation initiation factor IF- | 2 |
| QUC32_RS11715 |  | **2.69** |  | hypothetical protein- |  |
| QUC32_RS11725 |  | **2.64** |  | antitoxin Xre/MbcA/ParS toxin-binding domain-containing protein |  |
| QUC32_RS12745 |  | **2.33** | + | Phage shock protein operon transcriptional activator | 2 |
| QUC32_RS17275 |  | **2.50** | + | hypothetical protein | 2 |
| QUC32_RS18375 |  | **3.20** | + | hypothetical protein | 2 |
| QUC32_RS23085 |  | **2.26** | + | hypothetical protein | 2 |
| QUC32_RS23090 |  | 1.66 | + | Phage T7 capsid assembly protein |  |
| QUC32_RS23095 |  | 1.51 | + | Major capsid protein |  |
| QUC32_RS23100 |  | 1.78 | + | hypothetical protein |  |
| QUC32_RS28260 |  | **2.17** | - | Lrp/AsnC family transcriptional regulator | pHR1a |
| QUC32_RS28270 |  | 1.75 | - | creatininase family protein |  |
| QUC32_RS28275 |  | 1.55 | - | carbon-nitrogen hydrolase family |  |
| QUC32_RS28280 |  | 1.59 | - | M14 metallopeptidase family protein |  |
| QUC32_RS28285 |  | 1.73 | - | MFS transporter |  |
| QUC32_RS28290 |  | 1.80 | - | amidohydrolase |  |
| QUC32_RS28295 |  | 1.83 | - | IclR family transcriptional regulator C-terminal domain-containing protein |  |
