## Supplementary material for "An orphan LuxR regulator controls plant developmental reprogramming during interaction with *Novosphingobium resinovorum* HR1a": Supp Table 4

**Supplementary Table S4. Top 20 genes with the highest absolute loadings contributing to PC1 in the principal component analysis (PCA) of clover transcriptomes.** Genes are ranked according to their loading values. Functional annotations were assigned based on sequence similarity searches; E-value indicates the significance of the match, and score corresponds to the alignment score obtained for the best hit.

| gene | loading | evalue | score | Description |
| --- | --- | --- | --- | --- |
| P8452_13180 | -0.00762421 | 0 | 1285 | Pumilio homolog 6 |
| P8452_50686 | -0.00761405 | 5.64E-169 | 486 | Belongs to the UDP-glycosyltransferase family |
| P8452_47332 | -0.00760774 | 0 | 1054 | UPF0392 protein |
| P8452_45758 | -0.00760733 | 0 | 1128 | Protein NRT1 PTR FAMILY |
| P8452_77285 | -0.00760566 | 0 | 1147 | amine oxidase |
| P8452_56400\|RHM1 | -0.00760414 | 0 | 1324 | rhamnose biosynthetic enzyme |
| P8452_50673 | -0.00759913 | 0 | 884 | Belongs to the class-I pyridine nucleotide-disulfide oxidoreductase family |
| P8452_05970\|PLP2 | -0.00759746 | 2.08E-282 | 774 | Lipolytic acyl hydrolase (LAH) |
| P8452_62463 | -0.0075972 | 3.46E-121 | 353 | transcription factor |
| P8452_76900\|GDPD5 | -0.00759679 | 2.92E-229 | 636 | glycerophosphoryl diester phosphodiesterase |
| P8452_50834 | -0.00759472 | 6.69E-166 | 469 | oxidoreductase activity |
| P8452_50834 | -0.00759472 | 3.65E-208 | 578 | oxidoreductase activity |
| P8452_74795\|ALIS3 | -0.00759192 | 1.38E-230 | 636 | ALA-interacting subunit 3-like |
| P8452_21772\|RGLG2 | -0.00759062 | 1.35E-265 | 738 | E3 ubiquitin-protein ligase |
| P8452_12434 | -0.00758811 | 5.15E-228 | 631 | Removal of H(2)O(2), oxidation of toxic reductants, biosynthesis and degradation of lignin, suberization, auxin catabolism, response to environmental stresses such as wounding, pathogen attack and oxidative stress |
| P8452_65187\|FLA10 | 0.00758669 | 2.34E-214 | 602 | Fasciclin-like arabinogalactan protein |
| P8452_49036 | -0.0075854 | 0 | 1008 | Protein of unknown function (DUF2985) |
| P8452_29357\|RLK7 | -0.00758495 | 0 | 984 | Belongs to the protein kinase superfamily. Ser Thr protein kinase family |
| P8452_20383\|PBL2 | -0.00758321 | 9.01E-243 | 671 | Protein kinase 2B, chloroplastic-like |
| P8452_62500\|DOGT1 | -0.00758301 | 3.09E-284 | 785 | Belongs to the UDP-glycosyltransferase family |
