## Supplementary material for "An orphan LuxR regulator controls plant developmental reprogramming during interaction with *Novosphingobium resinovorum* HR1a": Supp Fig 1

### Slide 1
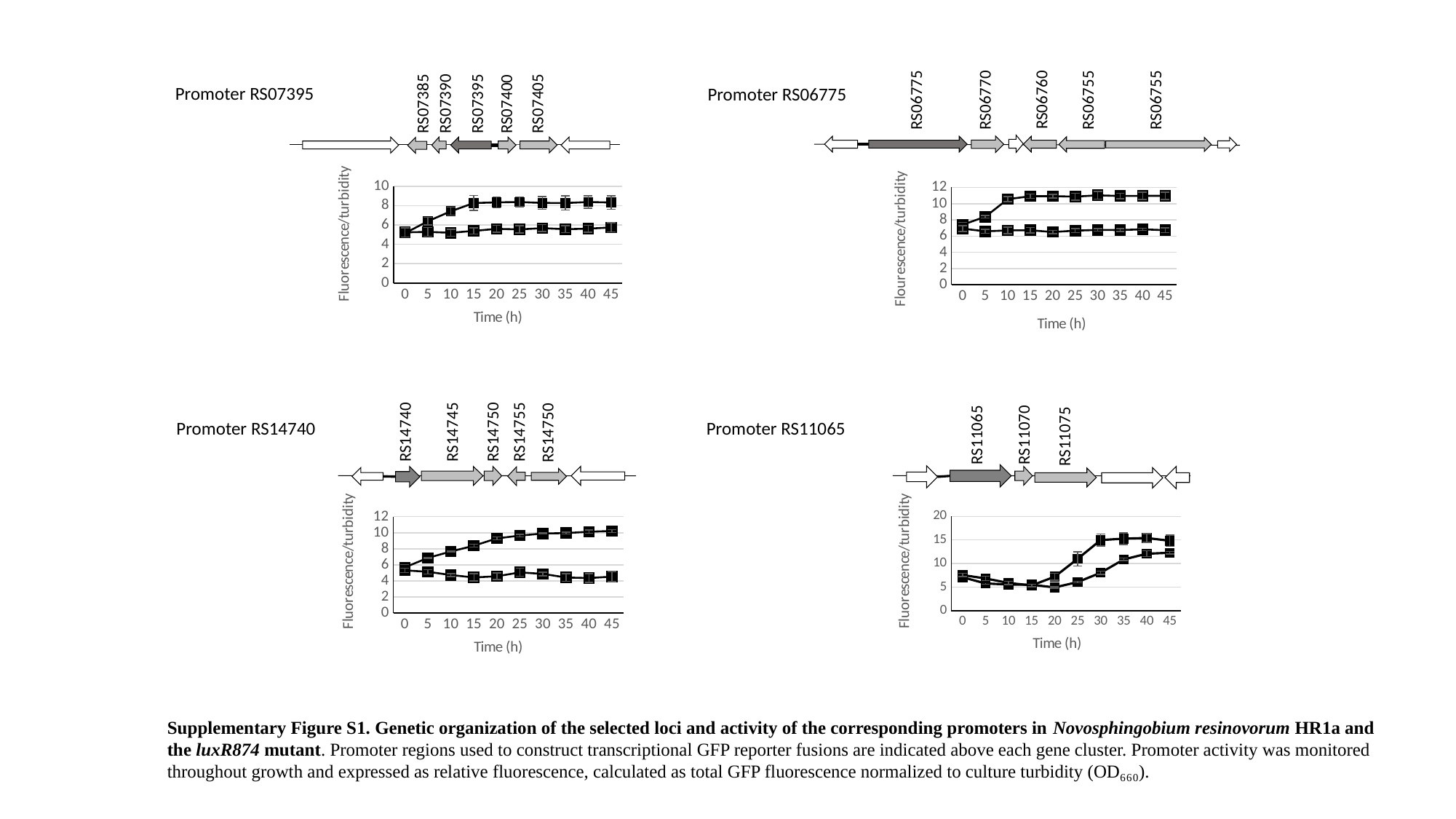

RS06760
RS06775
RS06770
RS06755
RS06755
RS07405
RS07395
RS07390
RS07385
RS07400
Promoter RS07395
Promoter RS06775
#### Chart
| Category | | |
|---|---|---|
| 0 | 7.405680119581465 | 6.919431988041853 |
| 5 | 8.385201793721974 | 6.586995515695068 |
| 10 | 10.556053811659194 | 6.7040358744394615 |
| 15 | 10.911808669656203 | 6.720777279521674 |
| 20 | 10.926756352765322 | 6.503437967115097 |
| 25 | 10.860986547085203 | 6.668011958146487 |
| 30 | 11.028400597907323 | 6.7605381165919285 |
| 35 | 10.941704035874439 | 6.744693572496263 |
| 40 | 10.961136023916291 | 6.833931240657697 |
| 45 | 10.968609865470853 | 6.731091180866966 |
#### Chart
| Category | | |
|---|---|---|
| 0 | 5.156881115969186 | 5.23776806162815 |
| 5 | 6.3907974182802425 | 5.2827399541952955 |
| 10 | 7.4172392254840736 | 5.193733083489486 |
| 15 | 8.265667291276285 | 5.386737455756819 |
| 20 | 8.33437434936498 | 5.601082656672912 |
| 25 | 8.373932958567563 | 5.541744742869041 |
| 30 | 8.277118467624401 | 5.672912762856549 |
| 35 | 8.266708307307933 | 5.566000416406413 |
| 40 | 8.370809910472621 | 5.6171143035602755 |
| 45 | 8.325005205080158 | 5.741515719342078 |RS11070
RS11065
RS11075
Promoter RS14740
Promoter RS11065
RS14745
RS14755
RS14750
RS14740
RS14750
#### Chart
| Category | HR1a | luxR874 |
|---|---|---|
| 0 | 7.08622537 | 7.55264208 |
| 5 | 5.77464195 | 6.8365079 |
| 10 | 5.53973374 | 5.87425257 |
| 15 | 5.50578934 | 5.36002993 |
| 20 | 4.92742885 | 7.19883395 |
| 25 | 6.06524582 | 11.0108966 |
| 30 | 8.04642929 | 14.9832169 |
| 35 | 10.8679397 | 15.2842907 |
| 40 | 12.0855441 | 15.4213103 |
| 45 | 12.2925415 | 14.8349433 |
#### Chart
| Category | | |
|---|---|---|
| 0 | 5.691422594142259 | 5.320316132031613 |
| 5 | 6.864132961413296 | 5.139237563923756 |
| 10 | 7.663877266387726 | 4.736169223616923 |
| 15 | 8.385402138540215 | 4.429102742910274 |
| 20 | 9.298000929800093 | 4.578219432821943 |
| 25 | 9.64344490934449 | 5.066364481636448 |
| 30 | 9.903881915388192 | 4.876569037656904 |
| 35 | 9.954556020455602 | 4.42747559274756 |
| 40 | 10.126336587633658 | 4.349837284983728 |
| 45 | 10.219665271966527 | 4.535564853556485 |Supplementary Figure S1. Genetic organization of the selected loci and activity of the corresponding promoters in Novosphingobium resinovorum HR1a and the luxR874 mutant. Promoter regions used to construct transcriptional GFP reporter fusions are indicated above each gene cluster. Promoter activity was monitored throughout growth and expressed as relative fluorescence, calculated as total GFP fluorescence normalized to culture turbidity (OD₆₆₀).
