## Supplementary material for "An orphan LuxR regulator controls plant developmental reprogramming during interaction with *Novosphingobium resinovorum* HR1a": Supp Fig 2

### Slide 1
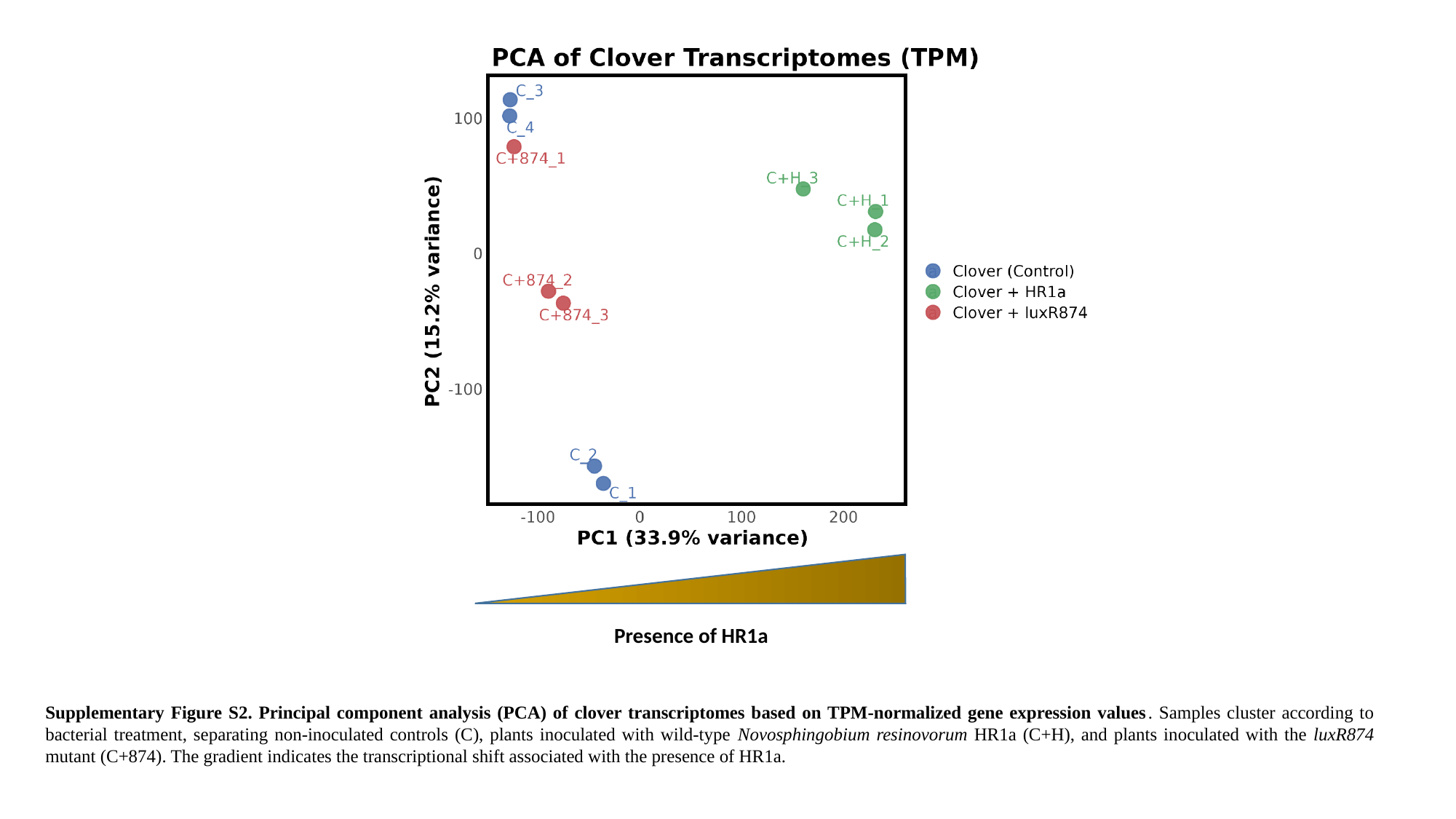

Presence of HR1a
Supplementary Figure S2. Principal component analysis (PCA) of clover transcriptomes based on TPM-normalized gene expression values. Samples cluster according to bacterial treatment, separating non-inoculated controls (C), plants inoculated with wild-type Novosphingobium resinovorum HR1a (C+H), and plants inoculated with the luxR874 mutant (C+874). The gradient indicates the transcriptional shift associated with the presence of HR1a.
