## Supplementary material for "An orphan LuxR regulator controls plant developmental reprogramming during interaction with *Novosphingobium resinovorum* HR1a": Supp. Fig 3

### Slide 1
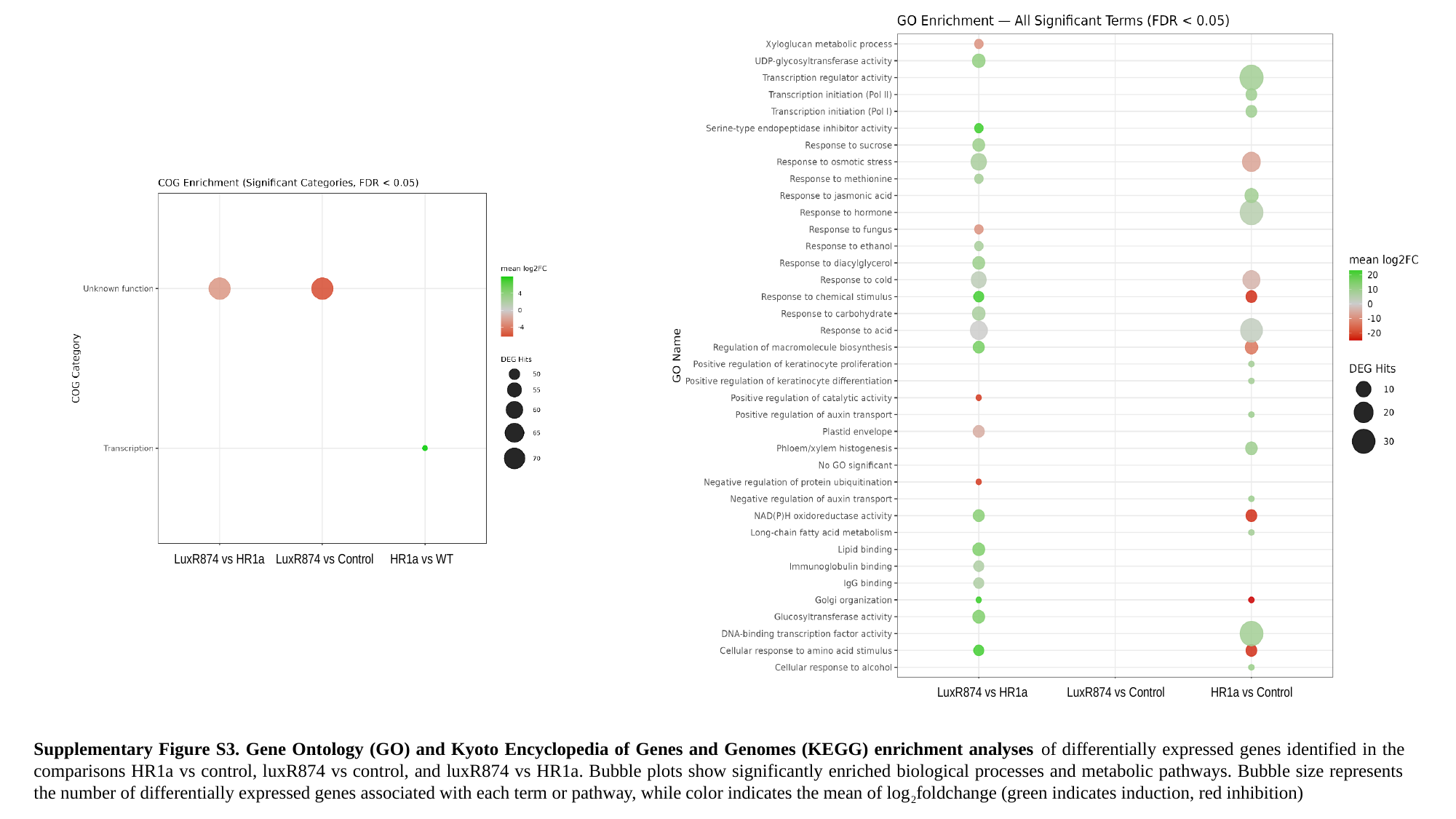

LuxR874 vs HR1a
LuxR874 vs Control
HR1a vs Control
LuxR874 vs HR1a
LuxR874 vs Control
HR1a vs WT
Supplementary Figure S3. Gene Ontology (GO) and Kyoto Encyclopedia of Genes and Genomes (KEGG) enrichment analyses of differentially expressed genes identified in the comparisons HR1a vs control, luxR874 vs control, and luxR874 vs HR1a. Bubble plots show significantly enriched biological processes and metabolic pathways. Bubble size represents the number of differentially expressed genes associated with each term or pathway, while color indicates the mean of log_2_foldchange (green indicates induction, red inhibition)
